# Cortical population codes for embedding sensory inputs into the prior context

**DOI:** 10.64898/2026.02.27.707941

**Authors:** Iacopo Hachen, Sebastian Reinartz, Alisea Stroligo, Alejandro Pequeno-Zurro, Mathew E. Diamond

## Abstract

According to theories of the brain as a predictive network, perceptual decisions result from integrating incoming sensory inputs with prior experience. A neuronal population implementing this form of computation must not only retain information about past events, but also combine this information with current sensory evidence. To examine how this integration occurs, we recorded extracellular activity from both primary sensory cortex and a target frontal region in rats performing stimulus categorization. Psychophysical analysis showed that judgments were history-dependent. Sensory cortex represented the current stimulus largely independently of prior stimuli and failed to account for the history-dependence seen in behavior. By contrast, frontal cortex embedded current input within a representation of prior sensory information through collinearity in coding dimensions. This mechanism – predominantly mediated by fast-spiking neurons – explained trial-to-trial variability in decisions. Our findings argue for distinct roles of cortical regions in predictive processing, and identify a frontal stage where current and prior sensory information converge to inform decisions.

## Introduction

Predictive accounts of perceptual decision-making entail the modulation of the current sensory information by recent context. In normative Bayesian frameworks, context is usually formalized as the “prior” (*1–3*). In perceptual decision-making, priors represent recently experienced sensory information (*4–11*), in addition to decision and reward contingencies (*12–19*). Previous work has revealed that perceptual dependencies unfold over much longer time spans than are traditionally attributed to low-level sensory adaptation (*5, 20, 21*): for instance, a current perceptual decision can be biased by a brief stimulus presented tens of seconds earlier, extending back over several trials (*5, 20*).

Three processes must underlie history-dependent perceptual decisions: (i) reliable encoding of current sensory input, (ii) storage of prior information, and (iii) merging of present sensory information and the prior to produce history-dependent choices. The third process constitutes the computation of a Bayesian “posterior,” an *a posteriori* estimate of the state of the world. Previous work has focused on identifying the substrate for sensory priors (*22*). Yet, to compute a posterior representation relevant for behavior, prior sensory history must converge with the neuronal code of the current input, an operation still poorly understood. Identifying the processing stage at which this computation occurs is crucial for understanding the operation of the brain as a predictive system. Even if representations of the priors of multiple task variables have been identified across widespread regions, including primary sensory cortex (*23–27*), these studies have largely characterized the prior in isolation. The critical step – merging the prior with new evidence – remains unsolved.

Here, we set out to identify the processing stage where traces of past stimuli influence current sensory representations. We trained rats to judge whisker vibrations as stronger or weaker than a fixed reference. As in earlier work (*5, 20*), a negative dependency between the current decision (*n*) and the preceding trial’s stimulus intensity (*n-1*) emerged: a strong past stimulus biased the next judgment toward “weak,” and vice versa. We recorded simultaneously from vibrissal primary somatosensory cortex (vS1) and vibrissal motor cortex (vM1). vS1 serves as the primary cortical relay for whisker kinematics, while vM1 – a main frontal target of vS1 (*28–32*) – contains whisker-encoding neurons (*28, 29*) and is involved in sensorimotor control (*33, 34*), orienting (*35–37*), delayed responding (*37, 38*), and perceptual decision-making (*39, 40, 29*). With extensive connectivity to sensory, parietal, and frontal areas (*28, 41, 36, 30*), vM1 is well positioned as a hub for context-dependent behavior.

If the negative aftereffect of stimulus *n-1* originates in primary sensory cortex or earlier pathways, it should be evident in vS1 activity. Alternatively, if trial-by-trial dependencies reflect higher-level contextual processing, history effects should appear more strongly in vM1 and covary with behavior, while being weaker and more decoupled in vS1.

Although phenomena of perceptual hysteresis such as adaptation have typically been characterized at the single-cell level, a complex computation such as merging longer-term priors and sensory inputs is likely to depend on population-wide mechanisms (*42, 43*). We therefore analyze how neuronal population codes represent present and past sensory inputs and how the intersection of these codes conditions history-dependent behavioral choices. We then identify the distinct roles of putative excitatory and inhibitory cell types in integrating present and past information. Finally, we describe a discrete transition in the population code for stimulus *n*-1: initially representing the ongoing sensory input, it shifts at reward collection into a memory-like representation that persists as the rat enters trial *n*. In sum, our findings characterize the neuronal code for a perceptual-decisional posterior that channels sensory history into behavior.

## Results

### Repulsive aftereffect of stimulus *n-1* on trial *n* choice

Each 500-ms vibration, delivered by a motor-driven plate (Figure 1A), was a sequence of speed values sampled from a half-Gaussian distribution (Methods). A single vibration was characterized by its nominal mean speed. Following earlier work (*40*), we define perceived intensity as the subjective experience of mean speed, and we refer to the task as intensity judgment. Nine intensities were used, each presented with a probability of 11.1% per trial, in randomized order (see Methods). An auditory cue, presented 400–600 ms after the vibration ended, instructed the rat to withdraw from the nose poke and choose between the two reward spouts. The left or right spout was baited depending on whether the trial’s vibration intensity was higher or lower than the category boundary. Correct choices were rewarded with a liquid delivery (pear juice diluted with water), accompanied by a reinforcing sound. For category-boundary stimuli, either choice was rewarded randomly (50% probability). For clarity, intensity is converted from mean speed (in mm/s) to a scale of −4 (lowest intensity) to 4 (highest intensity), with the reward rule boundary at 0. This linear-to-linear conversion does not affect the analyses.

**Fig. 1.**
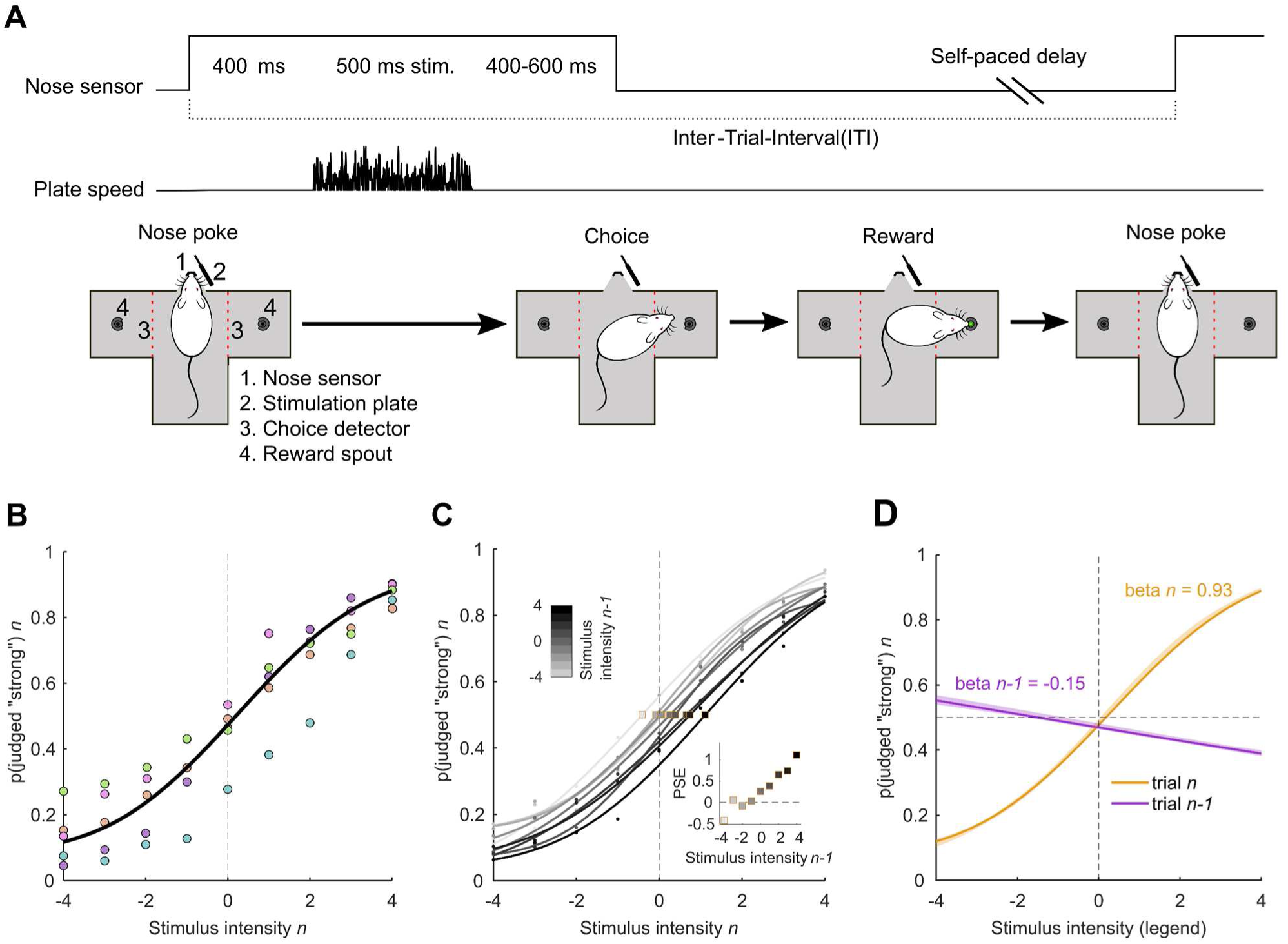
History-dependent perceptual decisions in rats during a vibrotactile psychophysical task. (**A**) Illustration of the task. By placing its snout in the nose poke, the rat triggered a whisker vibration. The rat withdrew upon the go cue to collect reward, and its turn direction was detected by an optic sensor (red dashed line). (**B**) Psychometric curve for trial *n*, describing the relation between stimulus intensity in trial *n* and the probability of judging stimulus *n* as “strong”, for all sessions in which neuronal data were recorded. Data of individual rats are color-coded dots. (**C**) Psychometric curves for trial *n*, conditioned on *n-1* intensity. The squares denote the Point of Subjective Equality (PSE) for each curve. The inset represents the PSE as a function of *n-1* stimulus intensity. (**D**) Predictions of multi-parameter psychometric models for the decision in trial *n*, as a function of stimulus intensity in both trials *n* and *n-1*. Shading represents 95% confidence intervals on model predictions, obtained with parametric bootstrapping.

Figure 1B shows the data of all five subjects (colored points) from sessions in which neuronal recordings were obtained. The psychometric curve (solid line) is the best cumulative Gaussian fit to the across-subject average. The curve resembles that seen in earlier studies (*5, 20*) (see Figure S1A for curves from a larger group of rats, including those not providing neuronal data). The repulsive influence of stimulus n-1 is evident in Figure 1C, where trial *n* psychometric curves are separated by *n-1* intensity. The inset displays the Point of Subjective Equality (PSE) for these curves, showing a linear dependence on stimulus *n-1* intensity (Pearson’s *r* = 0.97, *p* < 0.001). Further statistical verification of this repulsive effect is given in Figure 1D, where a multi-parameter psychometric model is fit on the dataset using as predictors not only intensity *n*, but additionally intensity *n-1* (see Methods). This fit yields weights for each predictor, which, through the logistic function, translate into choice probabilities. The resulting curves reveal a strong positive relation between trial *n* intensity and *n* choice (orange sigmoidal curve), and a negative relation between *n-1* intensity and *n* choice (violet), replicating the repulsive effect of Figure 1C. Control analyses (Figure S1B) confirm that this repulsive effect arises from history-dependent perceptual decision-making rather than unintended correlations or structures within stimulus sequences. Together, Figures 1C-D show that recent sensory history systematically biases perceptual judgments, giving rise to repulsive serial dependence in perceived intensity.

### Encoding of sensory inputs in vS1 and vM1 neurons

To establish how current and past sensory information is encoded in cortical populations – and how such encoding relates to history-dependent decision making – we recorded extracellular activity from neurons in vS1 and vM1 of rats performing the categorization task of Figure 1. Approximate locations of the multi-electrode array recordings are shown in the middle plot of Figure 2A (see Methods for implant coordinates). The dataset includes 264 units from vS1 and 594 units from vM1. A “unit” refers either to a clearly isolated single neuron or else a cluster formed of a small set of single-neurons (see Methods).

**Fig. 2.**
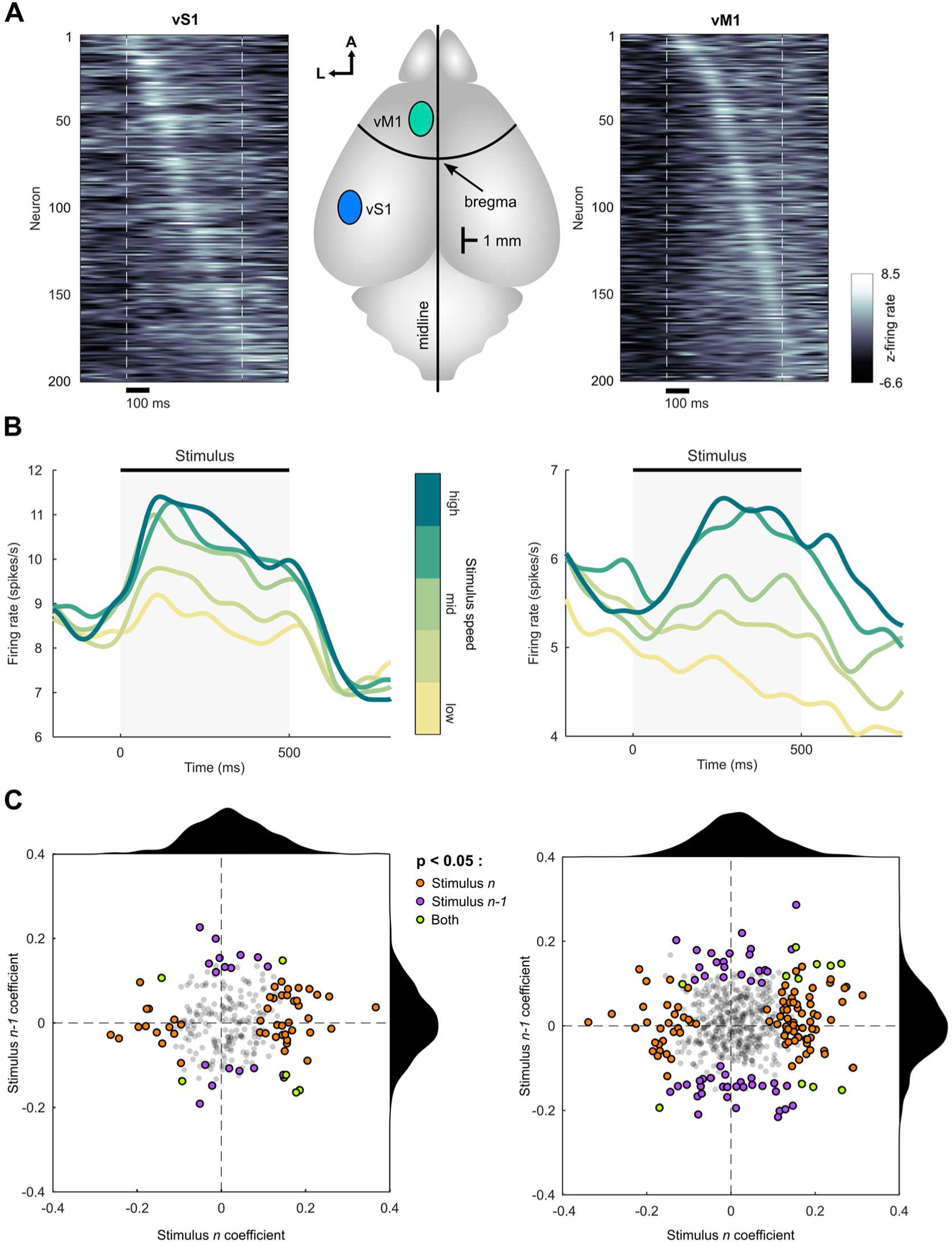
Individual-unit sensory responses in vS1 and vM1. (**A**) Left and right plots: z-scored responses for the 200 most stimulus-responsive units in vS1 and vM1, respectively. Units are further sorted by latency of peak firing rate, in sequential order. Dashed lines mark stimulus onset and offset. Center: locations of the recording sites in vS1 and vM1. (**B**) Left: average PSTH of vS1 neurons positively tuned to stimulus intensity, for 5 different intensity levels (grouped into 2 levels per stimulus category). The intensity value corresponding to the category boundary is labelled “mid.” Signals were smoothed with a forward-backward Gaussian filter (σ = 25 ms). The apparent response “leakage” – where firing rate increases prior to stimulus onset – results from the filtering procedure. Right: same analysis for vM1. (**C**) Distribution of coefficient values for stimulus *n* and stimulus *n-1*, in vS1 (left) and vM1 (right). Each dot represents a unit, colored according to its coding profile.

To identify stimulus-responsive units, we compared firing rates during the stimulus period (0–500 ms) to baseline activity (−200 to 0 ms). The 200 most responsive units from each area are shown in Figure 2A (left: vS1; right: vM1), sorted by peak response time. In vS1 a significant portion of the population is characterized by early responses to vibrissal stimulation whereas vM1 units display more gradual recruitment. For example, 68 vS1 units and 38 vM1 units (out of the selected 200) peaked within the first 200 ms of stimulation. A full distribution of peak times is provided in Figure S2.

We then focused on neurons modulated by stimulus intensity on the current trial (*n*), as these could directly contribute to intensity judgments. Intensity-tuned neurons were identified via Spearman’s correlation between stimulus intensity and firing rate during the stimulus window (0–500 ms), with a significance criterion of *p* < 0.05. In vS1, 54 of 264 units (20%) showed significant tuning, well above the 5% chance level (*p* < 0.001, binomial test). Of these, 38 were positively tuned and 16 negatively tuned. In vM1, 105 of 594 units (18%) were significantly tuned, also above chance (*p* < 0.001), with 71 positively and 33 negatively tuned.

Population-average PSTHs for the positively tuned neurons, with the 9 intensity values separated into 5 bins, further reveals differences in the response profiles between areas (Figure 2B). In vS1, response onset is steep; short-term adaptation can be observed in the decay of average firing rate starting about 100 ms after vibration onset and by the depression of firing rate below the pre-stimulus level following vibration offset. In vM1, firing rates are lower overall and response buildup is more gradual. Differently from vS1, at vibration offset firing rate decays gradually and thus continues to carry information until the end of the examined post-stimulus period.

Next, we looked for the effects of the previous trial (*n-1*) during the presentation of stimulus *n* by examining firing during trial *n* as a function of stimulus intensity on trial *n-1*. Because on trial *n* rats might have an innate or learned tendency to repeat or else to reverse the *n-1* choice (stick or switch, respectively), independently of the stimulus intensity of *n-1*, the real sensory-perceptual aftereffect of stimulus *n-1* could be confounded by post-perceptual decisional processes (*13, 14, 44*). To isolate sensory aftereffects from choice-related biases, we controlled for *n-1* choice by including in the analyzed pool of trials an equal number of trials where *n-1* choice was “strong” and “weak” (see Methods). With the potential effect of *n-1* choice thus cancelled, 22 of 264 vS1 units (8%) were significantly modulated by stimulus *n-1*, 12 positively and 10 negatively. This exceeds chance level (*p* = 0.0074, binomial test) but was significantly lower than the percent tuned to stimulus *n* (*p* < 0.001, Fisher’s exact test). The trial *n-1* correlation coefficients were significantly smaller in magnitude than those of the trial *n* coefficients (mean absolute values of 0.057 and 0.073, respectively; *p* = 0.0039, rank sum test). If the “memory” of stimulus *n-1* held by individual vS1 units were to condition the stimulus *n* response and consequently to give rise to the observed behavioral repulsion (Figure 1C-D), it would follow that the tuning to stimulus *n-1* would be systematically opposite to the tuning to stimulus *n*: the set of units most strongly tuned to stimulus *n* would overlap the set of units affected by stimulus *n-1*, but with opposite sign. Arguing against this account, Figure 2C, left, shows that only 6 units were significantly tuned to both *n* and *n-1* (4 with opposite sign, 2 with same sign), not different from the chance level of co-tuning as computed from the proportions of neurons tuned to the two trials separately (*p* = 0.129, binomial test).

In vM1, 63 of 594 units (11%) were significantly modulated by stimulus *n-1*, exceeding chance level (*p* < 0.001, binomial test), 34 positively and 30 negatively. As in vS1, this was significantly lower than the percent modulated by stimulus *n* (*p* < 0.001, Fisher’s exact test). Also as in vS1, the trial *n-1* correlation coefficients were significantly smaller in magnitude than those of the trial *n* coefficients (mean absolute values of 0.06 and 0.073, respectively; *p* = 0.0012, rank sum test). Overall, *n* versus *n-1* coding (quantified as the percent of all units encoding *n* divided by the percent of all units encoding *n-1*) did not significantly differ between vS1 and vM1 (*p* = 0.1937, Fisher’s exact test). As seen in Figure 2C right, only 11 vM1 neurons encoded both trials (4 with opposite sign, 7 with same sign), not different (*p* = 0.1118, binomial test) from the chance level expected if stimulus *n* and stimulus *n-1* coding were independent.

Overall, both vS1 and vM1 show a small but significant influence of stimulus *n-1* on trial *n* responses at the individual-unit level. However, rather than a single coding population showing sign-reversed tuning, *n* and *n-1* modulation appear to arise independently within groups of neurons.

### Population coding of history-dependent sensory processing in vS1 and vM1

In the preceding section, vS1 and vM1 units showed stimulus *n-1* coding that was independent of their stimulus *n* coding, suggesting that stimulus history effects, if processed in these regions, are embodied not at the single-unit level, but at the level of population dynamics. Populations can exhibit features that are not fully predictable from their individual components (*42, 43*). To test for such features, we trained decoders to classify stimulus *n* as “weak” or “strong” – mimicking the rat’s task – based on linear combinations of firing rates from all recorded neurons. Trials were grouped across sessions into pseudo-simultaneous population states (see Methods). We balanced the number of trials according to the category of stimuli *n-1* and *n*, where “weak” and “strong” categories include the four stimuli below and above the boundary stimulus, respectively. This yielded four distinct, equal-sized conditions of stimulus *n-1* to *n* transition (see Figure S3A). Trials where either stimulus *n-1* or *n* was at the category boundary were excluded. Since the decoders were trained only on stimulus *n*, the category of stimulus *n-1* was “hidden” to the classifier.

Figure 3A shows the average cross-validated decoding accuracy – color-coded for statistical significance – for vS1 and vM1 population activity in classifying stimulus *n*. The decoder was applied in a 400-ms sliding window across trial time. Only stimulus-*n*-tuned units were included in the figure (using all units yielded similar results; see Figures S3C-D). Accuracy is plotted at the center of each decoding window.

**Fig. 3.**
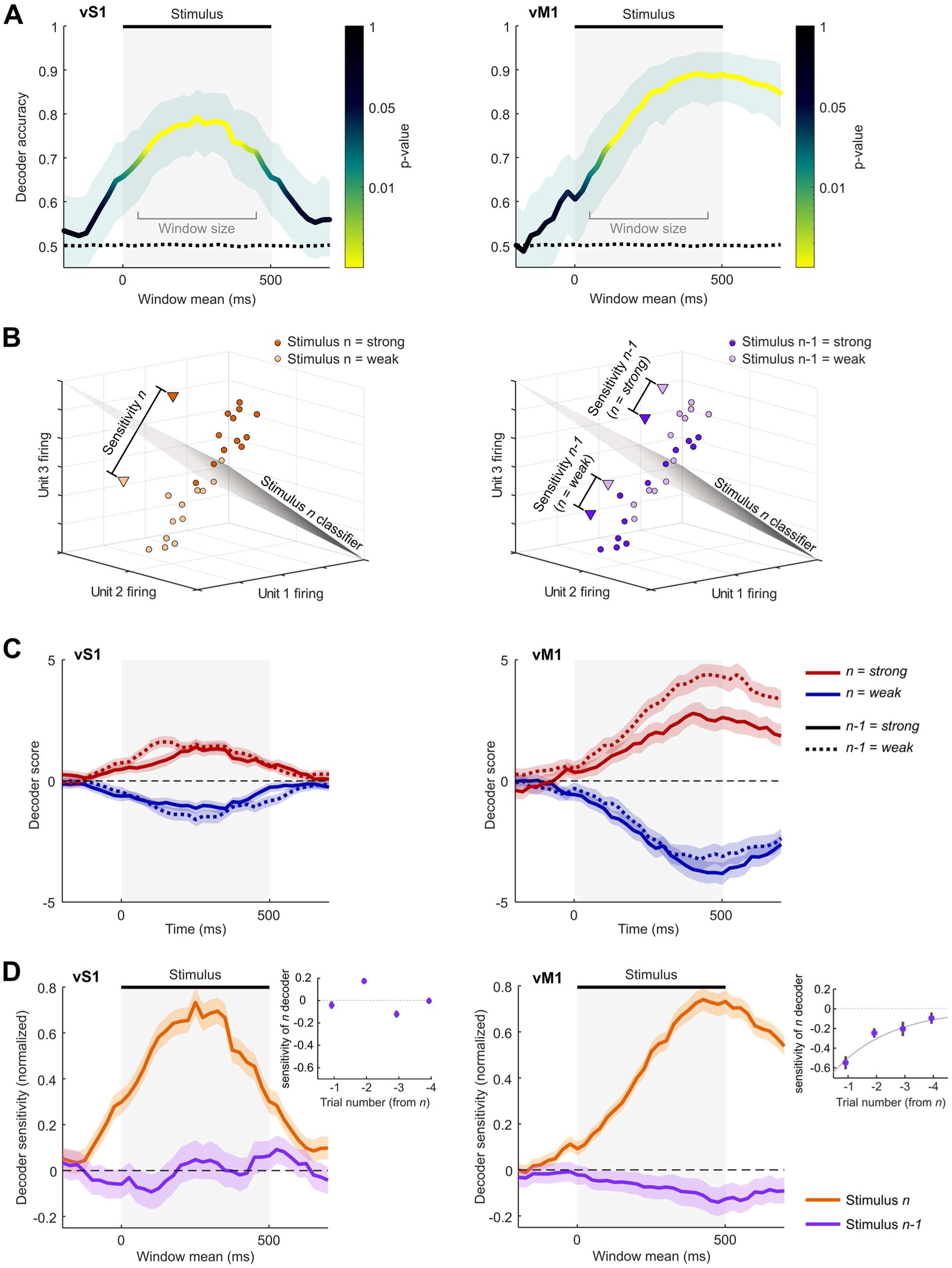
Population decoding of stimulus *n*. (**A**) Decoding accuracy of stimulus *n* from the firing rates of vS1 (left) and vM1 (right) populations in progressive time windows. The dotted line corresponds to decoding accuracy after randomly shuffling trial labels (Methods). (**B**) Schematic depiction of the preceding stimulus influence on decoding sensitivity. Left: all instances of trial *n* are considered independently of stimulus *n-1* category, with each point indicating the decoded stimulus category (strong above and to the right of the plane, weak below and to the left). Right: all instances of trial *n* are labeled according to stimulus *n-1* category. (**C**) Decoder scores, over time, of vS1 (left) and vM1 (right) populations on trial *n*, separated according to whether *n-1* was weak or strong. (**D**) Decoder sensitivity, over time, to stimulus *n* (orange) and to stimulus *n-1* (violet) of vS1 (left) and vM1 (right) populations on trial *n*. For comparability between different analyses, all sensitivity values were normalized by dividing by their maximum. All decoding analyses in the figure were run in 400 ms time windows, separated by 25 ms. Abscissae denote the central time point of the analysis windows. Shades represent 95% confidence intervals. The inset shows the effect of trials *n-1* to *n-4* on decoder sensitivity for vS1 and vM1. The sensitivity value is taken as the average of the time series between 0 and 500 ms and error bars represent SEM within time series.

In vS1 (left panel), the classifier performs significantly above chance for the entire stimulus period (the brief rise in accuracy before stimulus onset is due to the decoding window overlapping with the early stimulus response, extending 200 ms forward). Accuracy declines rapidly after stimulus offset. This temporal profile, including a peak at the center of the stimulus window, is consistent with a robust encoding of real-time sensory information in vS1 (*40, 45, 46*).

In vM1 (right panel), decoder accuracy increases gradually during the stimulus and peaks between 400–500 ms—when the decoding window encompassed both the end of the stimulus and the early post-stimulus period. Unlike vS1, the sustained accuracy in vM1 after stimulus offset persists longer than can be explained by window overlap, indicating that stimulus category information is conserved in vM1 even after the vibration has ended.

Although binary classification accuracy is informative about the reliability of signals carried by the neuronal populations, we expect stimulus encoding to occur in a graded manner. Therefore, we also examined the *distance* between the vector of firing rates in the population space and the decoder’s decision boundary—a continuous variable termed the decoder score (see Methods). Positive and negative scores indicate assignment to the strong and weak categories, respectively. Observations closer to the boundary are more ambiguous and are more likely to be assigned to one or the other category by chance, while those farther away are more reliable. Thus, by calculating the difference in average scores between trials with strong and weak stimuli, we can obtain a measure of decoder sensitivity – how separable the responses to weak and strong stimuli are on average.

Figure 3B schematizes this logic. The same set of trials is shown in both panels. On the left, decoding outcomes for trial *n* are shown irrespective of stimulus *n-1*. Each point, colored by true stimulus category, indicates the decoder output (strong above/right with respect to the plane, weak below/left); a few decoding errors can be seen – strong inputs lying on the “weak” side of the boundary, and vice versa. Decoder sensitivity is quantified as the distance between the mean positions (triangles) of trials from the two stimulus categories, strong and weak. On the right, the same trials are relabeled by the category of stimulus *n-1*. This allows us to assess the influence of the preceding trial by examining *n-1*-evoked shifts in the decoder score for stimulus *n*. If a weak *n-1* biases decoding toward “strong,” and a strong *n-1* biases it toward “weak,” regardless of whether *n* itself is strong or weak, as schematized here, the pattern would indicate a repulsive, history-dependent effect in population coding.

Although Figure 3B shows, for simplicity, a schematic for a single time window during stimulus presentation, this analysis can be extended across time. Figure 3C shows real decoder scores across trial time (400-ms windows). In vS1 (left panel), strong and weak stimulus *n* trials are shown separately (red and blue, respectively) and additionally split by strong (solid) and weak (dashed) *n-1* category. Decoder scores peak during the stimulus and are only minimally affected by stimulus *n-1*, suggesting a lack of history dependence. By contrast, vM1 decoder scores (right panel) rise more slowly and peak near the end of stimulus presentation. Notably, trials preceded by a weak *n-1* (dashed) consistently show scores that are shifted upwards with respect to trials preceded by a strong *n-1* (solid), whether *n* itself was weak or strong. This indicates that prior stimulus history influences decoding of the current trial. The *n-1* effect is more pronounced when stimulus *n* is strong.

Figure 3D quantifies decoder *sensitivity* over time – the difference in average scores between weak and strong trials. For vS1 (left plot), sensitivity to stimulus *n* (orange trace) tracks the decoder’s accuracy curve (Figure 3A, left), quickly rising during stimulation. Although the decoder was not trained to classify stimulus *n-1*, we used the balanced design to assess how the stimulus *n* output score depends on stimulus *n-1* category. We computed the difference in decoder scores for trial *n* depending on whether *n-1* was weak or strong (violet trace). If the repulsive bias observed in behavior were mediated by widespread adaptation of the vS1 population, this difference would be negative. Instead, the plot reveals no consistent effect during the stimulus period. Only a brief *positive* deviation appears near stimulus offset – more pronounced when all units (including untuned ones) are included (Figure S3 – possibly reflecting feedback signals from downstream areas.

For comparison, in vM1 (Figure 3D, right plot), decoder sensitivity to stimulus *n* (orange trace) mirrors the trend in classification accuracy (Figure 3A, right), rising steadily and remaining elevated after stimulation. As for vS1, we computed the difference in decoder scores for trial *n* depending on whether *n-1* was weak or strong (violet trace) and found a robust negative shift that develops during the stimulus and peaks at the end of the stimulus presentation, around the same time as the peak in stimulus *n* sensitivity. This significant and persistent negative value indicates that *n-1* systematically biases the decoding of *n* – in line with the repulsive behavioral bias observed psychophysically (Figure 1).

Previous studies using the same experimental paradigm demonstrated that repulsive perceptual effects arise not only from the immediately preceding stimulus (*n-1*) but extend over multiple trials (*5, 20*). To test for analogous history dependence in the cortical representations, we measured the average sensitivity of the trial-*n* population decoder to the stimulus of trials *n-1* to *n-4*. As shown in the Figure 3D insets, the effect is well described by an exponential decay trend in vM1 – where negative values indicate repulsion – confirming that history effects in vM1, as in behavior itself, extend beyond the immediately preceding trial. In vS1, no systematic effects are seen. In summary, the population-level representation of stimulus *n* in vM1, but not in vS1, is modulated by prior trial history. In the following section, we seek direct evidence for the contribution of vM1 to choice behavior.

### Role of vM1 population coding in decisional output

Comparing the temporal profiles of the vS1 and vM1 decoders reveals a key difference: while vS1 sensitivity declines after ∼250 ms, vM1 sensitivity continues to rise. The fact that better vM1 readout of the stimulus category can be achieved close to vibration offset, extending to early post-stimulus windows, can be interpreted in two ways: (i) vM1 accumulates information over time by integrating sensory input from vS1, and/or (ii) vM1 reflects processes related to choice formation and/or motor preparation. Distinguishing between these alternatives is non-trivial. In the dataset used for vM1 decoding, rats made correct choices on 78% of trials, meaning that two hypothetical representations – one of the stimulus category and one of the decision – would overlap 78% of the time, making it difficult to discern whether the sampled activity vectors correspond to stimulus category or choice.

Nonetheless, decoding of the stimulus from vM1 reached 89% accuracy (Figure 3A), exceeding behavioral performance by 11% (*p* < 0.001, signed-rank test on cross-validation folds; Figure S4). The fact that vM1 carries stimulus information that is not fully translated into choice suggests that its activity reflects a pre-decisional perceptual representation rather than a purely decisional code, which would collapse directly into the animal’s expressed choice. In vS1, decoder accuracy was 79%, significantly better than the behavioral accuracy of 75% on the same trials (*p* < 0.001; Figure S4). Interestingly, vM1 outperforms vS1 in encoding the stimulus category (Figure 3 and Figure S3).

To further assess how vS1 and vM1 representations are converted to behavioral output, we repeated the decoding analyses by using the rat’s choice as the trial *n* label instead of stimulus category. In vS1, choice decoding was less accurate than stimulus category decoding (Figure S4). In contrast, vM1 showed similar or slightly better performance when decoding choice compared to stimulus category (Figure S4). This supports a dual role for vM1: while its activity reflects sensory-perceptual information, it also participates in choice formation, bridging perceptual representation and motor output.

Shared variability between the decoder output and choice would support the idea that the history dependence seen in the vM1 representation contributes to behavior (Figure 4A). Evidence for such shared variability is shown in Figure 4B, where each point reflects the average decoder score (abscissa) and probability of a “strong” behavioral choice (ordinate) for the four trial sequence conditions. Following strong *n-1* trials, the decoder’s output is shifted toward “weak,” just as rats were more likely to judge stimulus *n* as weak under the same conditions. The opposite pattern holds following weak *n-1* trials.

**Fig. 4.**
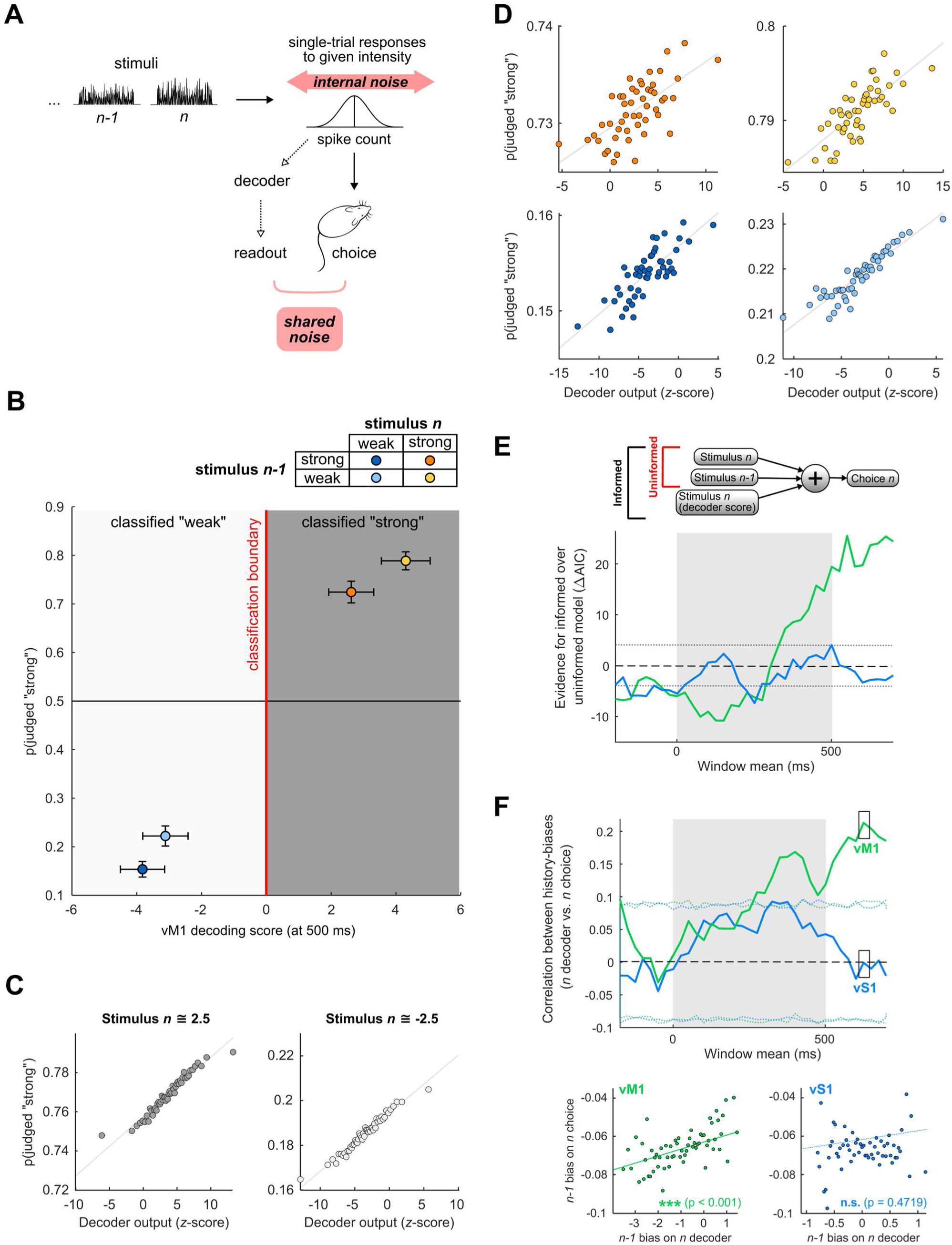
Relation between decoding scores and perceptual decisions. (**A**) Scheme of the influence of endogenous variability on decoding scores. If a neuronal representation of sensory information is responsible for guiding the behavioral choice, variability in the choice should be partly explained by variability in the decoded representation. (**B**) Average cross-validated decoding scores and average rat judgments across 500 samples. Color code table refers to the four different *n-1*-to-*n* sequences. Error bars in the main plots represent standard deviations. (**C**) Average probability of “strong” judgment for each of the stimulus *n* categories as a function of decoding score for stimulus *n*. Only for visualization, scores were discretized in 50 equipopulated bins. To avoid a trivial correlation due to a wide span of intensity values, only trials in which the mean intensity value was 2.5+-0.1 (left) or −2.5+-0.1 (right) were selected (6-7% of the trials). Gray lines are regression slopes. Pearson’s *ρ* (in non-binned data) = 0.1907 (left) and 0.1535 (right); both *p*-values < 0.001. (**D**) Average probability of “strong” judgment for each of the four possible trial sequences represented in (B) as a function of decoding score for stimulus *n* (color code as (B)). Only for visualization, scores were discretized in 50 equally-populated bins. As for (C), only trials in which the mean intensity value was 2.5+-0.1 (left) or −2.5+-0.1 (right) were selected. Gray lines are regression slopes. From top left, clockwise, Pearson’s *ρ* (in non-binned data) = 0.04, 0.05, 0.06, 0.11; all *p*-values < 0.001. (**E**) Upper: schematic of the linear regression used to predict choice as a function of task variables and decoding score for stimulus *n*. Lower: improvement of the vS1 (blue) and vM1 (green) informed models over the respective uninformed models in predicting rat choices, measured as difference in AIC, for all time windows considered in decoding analyses (upper and lower dashed lines correspond to a difference of 5 AIC units). (**F**) Upper: Pearson correlation between the history-dependent bias in the rat choices and the bias in the stimulus *n* representation decoded from vM1. *p*-values were extracted by comparing the relevant statistic with a distribution obtained after 1,000 random permutations of the data, in each time window (lines correspond to 95% range). Lower: relation between bias in the rat choices and bias in the stimulus *n* representation at the timepoint given by the box in the upper plot, for both vS1 and vM1.

A second form of shared variability is of interest: if vM1 has a causal role in transforming the trial *n* sensory input into a decisional output, the trial-to-trial fluctuations in the vM1 decoder scores of stimulus *n* should predict trial-to-trial fluctuations in behavior (*47–49*). Testing this begins with the reasoning that, for a given stimulus category (e.g., strong), randomly selected trial subsets will vary in both decoder scores and choice probabilities. If vM1 activity influences behavior, then in groups of trials where the stimulus was in the strong category and the rats made the “strong” judgment more frequently than in the overall mean, the population state should also be shifted towards “strong” with respect to the mean; the opposite effect should be seen for trials selected from the weak stimulus category.

Had we considered the full range of vibration intensity values, the covariance between score and choice would be trivial. Thus, to distinguish choice-related variability from stimulus-driven variability, we restricted the analysis to a narrow range of input intensities (see Methods). As shown in Figure 4C (left: strong trials, right: weak trials), there is a significant positive correlation between decoder scores and choice probability in both stimulus categories, indicating that trial-to-trial fluctuations in vM1 population activity are congruous with trial-to-trial fluctuations in judgment. A correlation between population activity and judgment, albeit weaker, was also found in vS1 (Figure S4).

Next, we asked whether the effect of stimulus *n-1* is mediated through the vM1 population state. If it is, then the correlation between decoder score and choice should persist even when controlling for the *n-1*-to-*n* sequence. That is, the correlation between the decoder’s scores and the rats’ choices must be seen independently of *n-1*, as if *n-1* ultimately acts through the vM1 population activity on trial *n*. To test this, we again split the data by trial *n* category (strong, weak) but further split the data based on stimulus *n-1* (strong, weak). As shown in Figure 4D, the positive correlation between decoder output and behavior remains robust across all four [*n-1, n*] combinations, supporting the idea that the influence of *n-1* on choice is reflected in – and exerted through – the vM1 representation of stimulus *n*.

To generalize the relationship between cortical population activity and behavioral output across the entire trial sequence, we employed a nested regression approach using two models to predict the probability of a “strong” choice (Figure 4E, top). The first model, termed “uninformed,” included only the raw intensity values (not labeled as category) of stimulus *n* and stimulus *n-1* as predictors. The second model, “informed,” added a third predictor: the stimulus *n* decoder score derived from the vM1 or vS1 population activity. Decoder scores were computed in sliding time windows (400 ms), and each iteration of the model used the score from the corresponding window. We adopted the Akaike Information Criterion (AIC) to compare the explanatory power of the models (see Methods). Figure 4E (lower plot) shows the differences in vS1 and vM1, uninformed minus informed (ΔAIC); any difference >5 (see dashed lines) is typically accepted as warranting the more complex model(*50*). The vM1-informed model (green) consistently outperforms the vM1-uninformed model. The time course of ΔAIC reveals that vM1 activity begins to enhance choice prediction around 200 ms after stimulus onset, with increasing predictive strength through the end of the stimulus and into the post-stimulus period, just before the “go” cue.

Applying the same analysis to vS1 activity shows no systematic advantage for the vS1-informed model over the vS1-uninformed model (Figure 4E, blue line). The negligible advantage afforded by the vS1 scores, consistent with the data of Figure S4, agrees with earlier results showing only a small contribution of vS1 neurons to behavioral reports (*51*). Coefficient estimates for the vM1 population-informed model are shown in Figure S4.

We next asked to what extent the negative bias imposed by stimulus *n-1* on the vM1 decoder could account for the behavioral bias observed in the rats’ choices. Across 500 cross-validation samples (each with 4,000 pseudo-trials), there was variation in the strength of both the decoder’s *n-1* sensitivity and the rats’ *n-1*-evoked choice bias – in a given 4,000-trial sample, *n-1* will exert more or less influence on trial *n* choice with respect to the average and the decoder’s *n-1* sensitivity will also vary with respect to the average. If the two measures covary, this would suggest a causal role for vM1 in mediating history-dependency. We compared the decoder/choice correlation of vM1 with that of vS1. For vS1 (Figure 4F, blue), the correlation between decoder bias and behavioral bias is weak and does not exceed chance level (*p* > 0.05; permutation test, dashed lines). In contrast, the correlation for vM1 (Figure 4F, green) is positive, highly significant, and especially prominent during the second half of the stimulus and into the post-stimulus period.

In summary, vM1 encodes the current stimulus (*n*) in a history-dependent manner (Figure 3C–D); additionally, vM1 trial-to-trial fluctuations – both in the representation of *n* (decoder score) and in the influence of *n-1* (negative sensitivity) – are significantly correlated with fluctuations in the rats’ choices (Figure 4). By contrast, vS1 exhibits minor history-dependence (Figure 3C–D). Further, the vS1 variability in stimulus *n* representation, and in the bias imposed by stimulus *n-1*, is not tightly linked to behavioral outcomes (Figure 4). These results argue against the thesis that the repulsive effect of prior stimuli is present at earliest sensory processing levels and relayed forward from there. Rather, they suggest that vS1 firing is more directly connected with the physical features of the current input while vM1 lies at the interface between stimulus representation and decision formation.

### Impact of stimulus *n–1* representation on trial *n* choice

If vM1 is one locus where history-dependent modulation occurs, then it must contain explicit information about stimulus *n-1*. We addressed this by training the classifier to decode stimulus *n–1* rather than the current stimulus (*n*), based on linear combinations of firing rates in vS1 or vM1. All neurons that met standard physiological selection criteria were included, regardless of their individual stimulus tuning (see Methods). To ensure that decoding pertained to stimulus *n–1* and not correlated variables such as previous motor action or decision, we selected a balanced set of trials – equal numbers of “strong” and “weak” choices – for each category of stimulus *n–1*.

In vS1 (Figure 5A, left), stimulus *n-1* can be decoded during trial *n* with only marginally significant accuracy (*p* < 0.05). This reinforces earlier evidence (Figure 3C-D) that vS1 plays a limited role in carrying forward information from previous trials into the current perceptual judgment. Notably, the signal is detectable only before the onset of stimulus *n*, weakens during the stimulus presentation, and disappears entirely before the behavioral response. In contrast, accuracy in decoding stimulus *n-1* from vM1 activity is significantly above-chance (*p* < 0.05) throughout trial *n*, with a peak during stimulus delivery (Figure 5A, right). Strikingly, in vM1, the highest *n-1* decoding accuracy occurs during the first half of stimulus *n* – precisely when the repulsive influence of *n-1* on the representation of *n* is strongest (Figure S3). Although the signal weakens after the stimulus offset, it remains detectable for the remainder of the examined window.

**Fig. 5.**
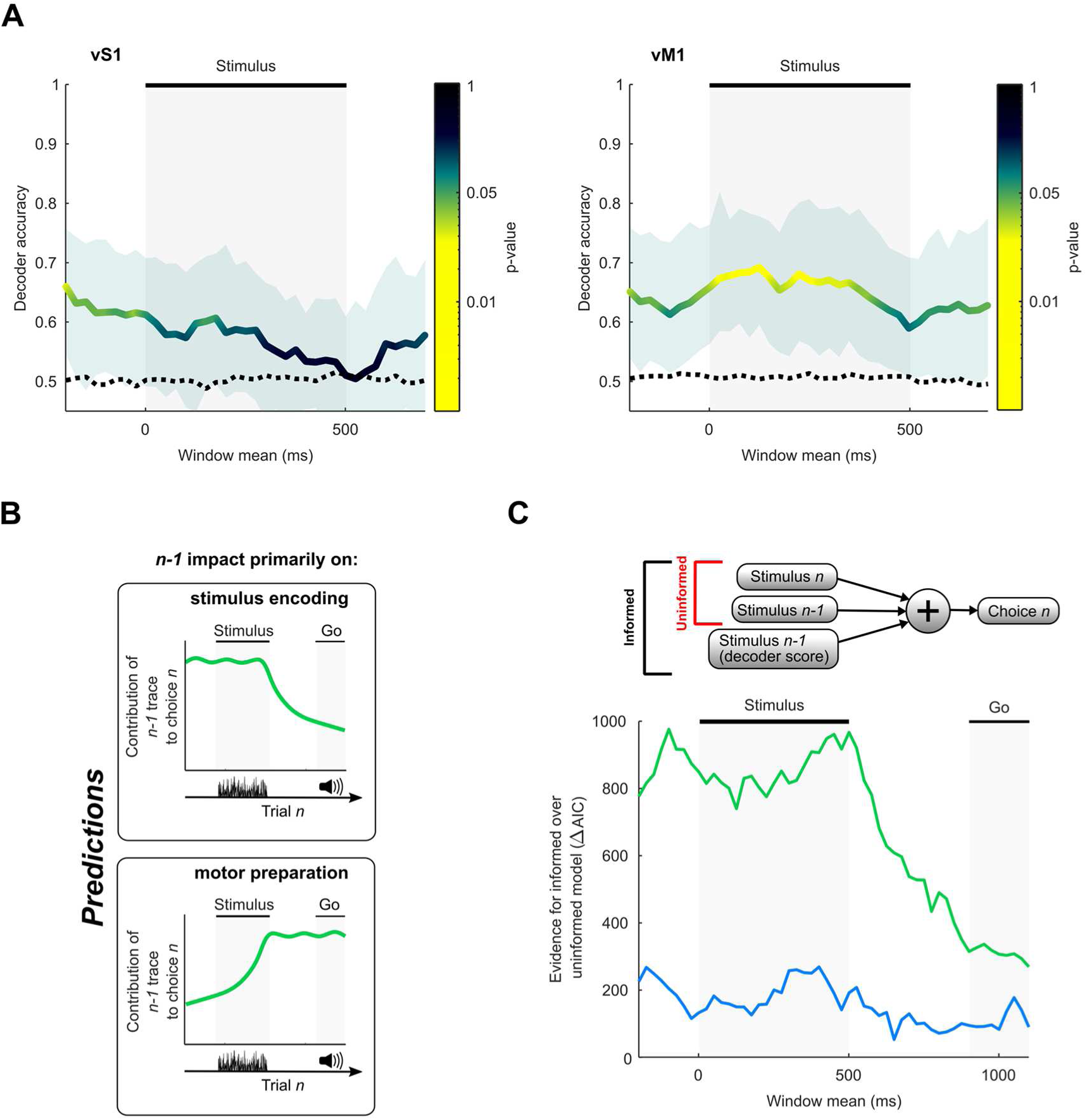
Population decoding of stimulus *n-1* and its relevance to behavioral choices. (**A**) Decoding accuracy of stimulus *n-1* from the firing rates of vS1 units (left) and vM1 units (right). The dotted lines represent the chance level obtained by randomly shuffling trial labels (see Methods). Analyses were run in 400 ms time windows, separated by 25 ms. Abscissae denote the central time point of the analysis windows. Shading represents 95% confidence intervals. (**B**) Predictions for two alternative mechanisms mediating the impact of stimulus *n-1* on choice *n* in vM1. (**C**) Upper: schematic of the linear regression used to predict choice as a function of task variables and decoding score for stimulus *n-1*. Lower: AIC difference-based improvement of the vS1 (blue) and vM1 (green) informed model over the uninformed model.

While the evidence so far implicates vM1 in history-dependent modulation, the computational role of this signal remains unclear: precisely how does stimulus *n-1* influence choice on trial *n*? Two scenarios are possible (Figure 5B). In the first, vM1 contributes to sensory-perceptual processing: as stimulus *n* is encoded, vM1 forms a history-dependent representation that other brain regions later transform into a motor plan. In the second, vM1 constitutes a later stage of the sensorimotor transformation, where the *n-1*-evoked bias directly drives motor planning or action selection. Rats may evaluate their decision even after stimulus *n* ends, until the go-cue prompts a response. If history-related activity predicts choice during stimulus encoding but not later, this would support a perceptual role; if the influence emerges near response initiation, it would point to action planning. To distinguish these alternatives, we asked when in the trial timeline history signals in vS1 and vM1 contribute most to predicting the upcoming choice.

As in Figure 4, we exploited trial-to-trial variability to link neuronal signals to behavior. Like any feature encoded in neuronal activity, the *n-1* representation varies across trials. If this variability contributes to behavioral choice, a model that includes the neuronal representation of *n-1*, in addition to the physical value of stimulus *n-1*, should better predict trial *n* choices than a model that includes only the physical value of stimulus *n-1*. Accordingly, we compared two linear models of choice behavior. The “uninformed” model included only the physical values of stimulus *n* and *n-1* as predictors. The “informed” model added the decoder score for stimulus *n-1*, derived from either vS1 or vM1 population activity.

We again used the AIC to compare the statistical explanatory power of the models. As in earlier analyses, we computed ΔAIC as the difference between uninformed and informed models; values greater than 5 (dashed line) indicate a substantial gain in explanatory power. As shown in Figure 5C, both the vS1 and the vM1 decoder scores provide measurable benefits in the prediction of trial *n* choices, although the advantage provided by the vS1 score (blue line) score is far below that provided by the vM1 score (green line). The predictive advantage provided by including the vM1 score declines sharply after the stimulus ends, suggesting that, even if the *n-1* signal persists in vM1 (Figure 5A), its relevance for decision-making wanes quickly – well before the initiation of motor output (i.e., the “go” cue). This temporal profile mirrors the scenario proposed in the upper panel of Figure 5B, suggesting that the *n-1* information carried by vM1 contributes primarily to the perceptual representation of stimulus *n*, rather than to motor preparation. Coefficient estimates for the vM1 population-informed model are shown in Figure S5.

It may be noted from the plot ordinates (Figure 4E) that the improvement achieved by including the population score of both vS1 and vM1 is much greater when employing the *n-1* score than when employing the *n* score. This can be explained by the fact that the *n* population score is highly collinear with stimulus *n*, providing information that is redundant, since the decoder reaches a much higher accuracy when decoding stimulus *n* (Figure 3A) than when decoding stimulus *n-1* (Figure 5A). When decoding *n-1,* the variability in the neuronal representation is less collinear with the ongoing stimulus, providing a greater amount of “new” information about the choice.

The regression coefficients of the vM1 informed model (Figure S5) are consistent with the repulsive effect of *n-1*. Specifically, the coefficient for the decoded trace of *n-1* is systematically negative, especially in vM1. This means that a positively biased representation of *n-1* (*n-1* stored as “stronger” than average) leads to a more negative judgment (“weaker”) of *n*, and vice versa. The picture emerges that the trace of *n-1* in vM1 significantly biases the coding of stimulus *n* (Figure 3D) and ultimately drives the repulsive bias in perceptual decisions on trial *n*.

### Cell-type specificity of vM1 population codes for stimulus *n* and *n-1*

Having established that vM1 mediates the convergence of present and past sensory information into behavioral choices, we next examined the features of the neuronal representation that might mediate such convergence. Linear decoders operate by multiplying each unit’s firing rate by a learned weight to classify stimuli. These weights form what we refer to as the *population code*, that is, the weights’ distribution that best discriminates stimuli across the classifier boundary. Figure 6A shows the decoding weights assigned to each of the 594 vM1 units for optimal discrimination of stimulus *n* (orange) and stimulus *n-1* (violet), ordered by increasing weight for *n*. The slight but significant negative slope of the linear fit to the *n-1* weights (violet line) hints that negative *n* weight may be weakly associated with positive *n-1* weights, and vice versa (this anti-correlation will be examined subsequently). Weights were averaged over time windows from 1 to 500 ms, as they were found to remain stable throughout the delivery of stimulus *n* (Figure S6). Notably, the code for *n* shows near-zero correlation with pre-stimulus windows (Figure 6B) – to be expected, as neuronal firing could not predict the upcoming stimulus – whereas the *n-1* code shows substantial correlation, suggesting that the code for *n-1* is present prior to *n* onset (Figure S6). This temporal overlap is consistent with a mechanism where the residual activity from *n-1* influences the encoding of *n*.

**Fig. 6.**
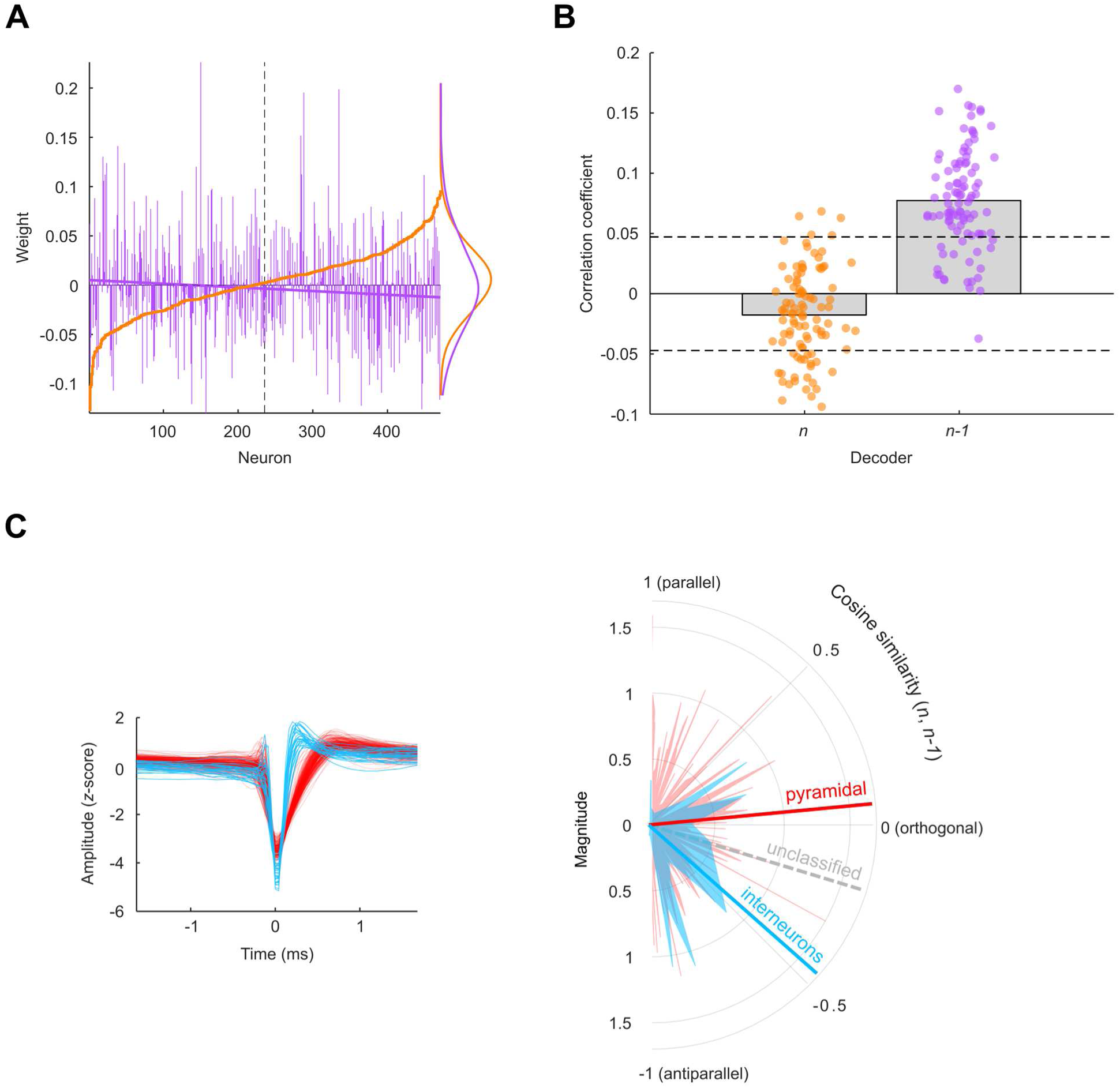
Distribution of population codes in vM1. (**A**) Neuronal weights assigned by the decoders for stimulus *n* (orange) and stimulus *n-1* (violet), sorted in increasing order relative to the stimulus *n* code. To maximize population size, all units meeting basic selection criteria (see Methods) were included in decoding regardless of stimulus tuning. Decoding accuracy for stimulus *n* was largely unaffected by population size in vM1 (Figure S3). (**B**) Correlation between the population code at the center of vibration delivery (250 ms) and the code at −200 ms (pre-stimulus), for the *n* and the *n-1* decoder. Individual points correspond to different resampled trial sets (see Methods). Dashed lines indicate the 95% range of a null distribution obtained through permutation test. (**C**) Left: average waveforms of units classified as fast-spiking (blue) and regular-spiking cells (red). Right: cosine similarity between decoding weights for stimulus *n* and *n-1*, computed separately for populations of regular-spiking neurons and fast-spiking. Polar plots show the cosine angle between *n* and *n-1* weights for each neuron (angular axis) and the magnitude of each unit’s 2D weight vector (radial axis). The angular axis is uniformly scaled for visualization.

The non-transposability between the *n-1* and *n* population codes in vM1 (Figure 6A) implies that during the presentation of stimulus *n*, two population codes coexist: one encoding *n*, the other encoding *n-1*. When *n-1* was strong, stimulus *n* tended to be encoded as weaker, and vice versa, suggesting a *repulsive interaction* between the two codes. This behavioral and neuronal pattern implies partial redundancy or overlap in the way *n* and *n-1* are represented – specifically, their population vectors point in *opposite* directions.

To explore this relationship more rigorously, we computed the cosine similarity between the decoding weight vectors for stimulus *n* (denoted *w*_*n*_) and stimulus *n-1* (*w*_*n*−1_). A cosine similarity of 0 would indicate orthogonality (no overlap), while a significantly negative value would confirm that *w*_*n*_ and *w*_*n*−1_ point in opposing directions. A positive similarity would contradict the repulsive interaction seen in earlier evidence. The observed cosine similarity was found to be significantly negative (*r* = −0.099; *p* = 0.0216, permutation test), confirming that the codes for *n* and *n-1* have opposing orientations.

Despite the low prevalence of individual units tuned to both *n* and *n-1* (Figure 2C), we hypothesize that subpopulations of cells, linearly combined, could have a role in mediating the repulsive effect of stimulus *n-1* on the stimulus *n* representation. Interneuron populations are a good candidate for mediating trial-by-trial context-dependency (*52–54*), including stimulus history (*55, 56*). We therefore identified putative pyramidal neurons and interneurons among the 594 units, using extracellular spike waveform features (specifically, peak-to-trough width (*56–58*); Figure 6C left). Of 173 classified units, 154 were labeled as pyramidal neurons and 19 as interneurons (interneuron-to-pyramidal ratio: 0.124). Within each cell class, we computed the cosine similarity between *w*_*n*_ and *w*_*n*−1_ (Figure 6C, right). Among putative pyramidal neurons, the cosine similarity was not significantly different from zero (similarity = 0.06; *p* = 0.21), indicating orthogonality – i.e., pyramidal cells tend to code for either *n* or *n-1*, but not both. In contrast, among interneurons the cosine similarity was significantly negative (−0.49; *p* = 0.022), suggesting that these neurons systematically flip their tuning between stimuli *n* and *n-1*. In the set of unclassified units, *w*_*n*_ and *w*_*n*−1_also retained a negative relationship (cosine similarity = −0.188; Figure 6C, dashed gray line), suggesting a mixture of both cell types within that cell population.

To assess whether this pronounced negative similarity in interneurons could occur by chance in equivalently sized samples of pyramidal neurons, we performed 20,000 random subsamplings of 19 pyramidal neurons each and computed the cosine similarity of their *w*_*n*_ and *w*_*n*−1_vectors. As shown in Figure S6, cosine values equal to or more negative than that of the interneurons were rare (*p* = 0.008). This control implies that the small set of interneurons contributed disproportionately to the overall negative similarity observed at the complete population level (–0.099).

### Episode-linked shift in cortical representations of past stimuli

In vM1, the representation of stimulus *n* is systematically influenced by the preceding stimulus (*n-1*) (Figure 3D). Indeed, *n-1* remains decodable throughout the presentation of *n* (Figure 5B). However, information about the current and preceding stimuli is carried by distinct population codes (Figure 6A). When the coding of *n-1* transitions from “real time” (during trial *n-1*) to “memory” (during trial *n*), the corresponding decision boundary must also shift. As schematized in Figure 7A, on the left a classifier (orange line) trained on population activity during *n-1* separates strong from weak inputs (filled vs. unfilled points). In the center, as the population configuration rotates counterclockwise (arrow; previous distribution in gray), the previously optimal boundary (orange) no longer distinguishes strong and weak responses. On the right, retraining identifies a new decision boundary (purple) that correctly classifies the memory representation of *n-1*.

**Figure 7.**
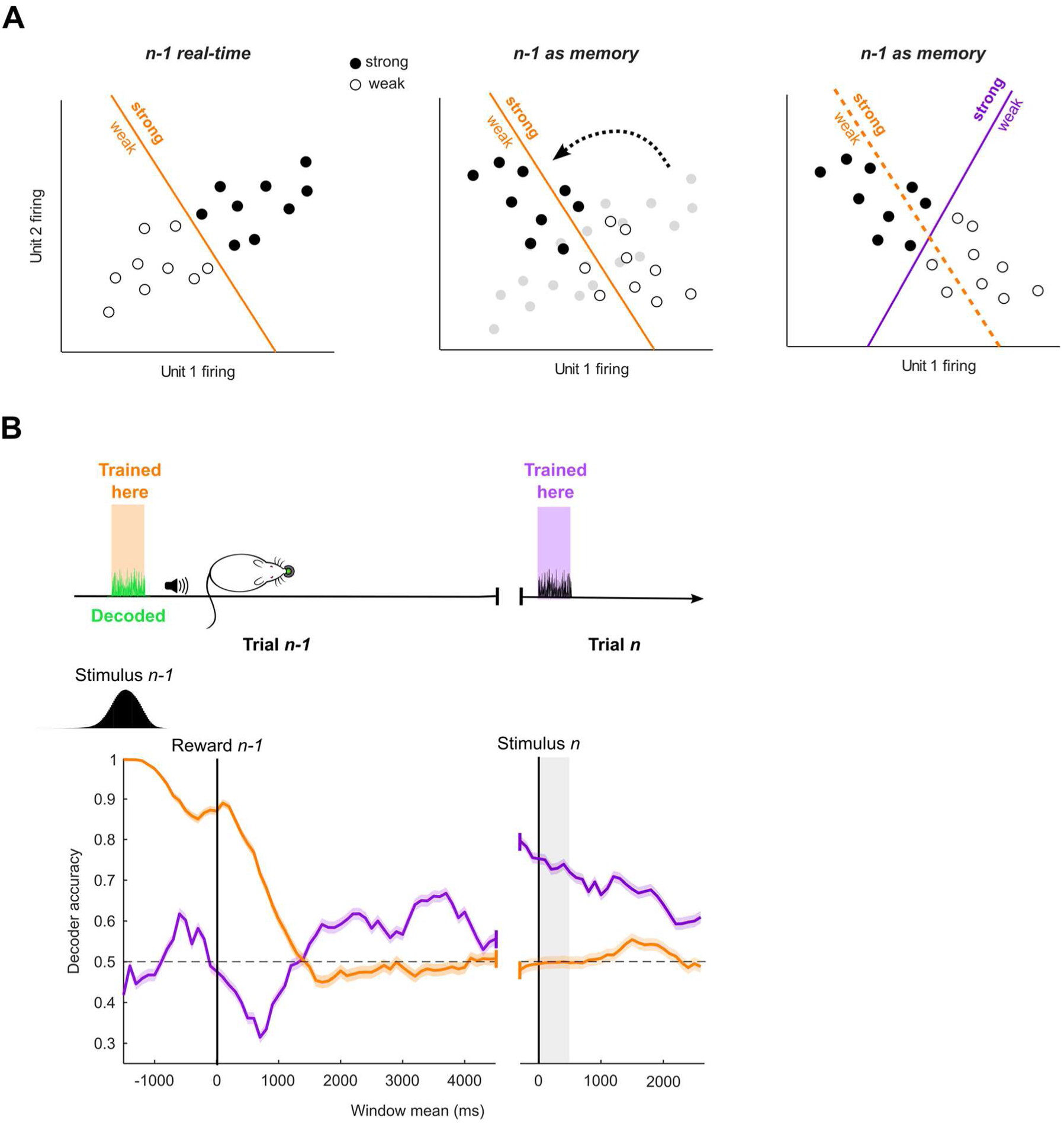
Transition between “real time” representation to “memory trace”. (**A**) Schematic to illustrate the rotation of the vM1 population codes in the transition from *n-1* to *n*. (**B**) Upper: schematic of the cross-temporal validation of decoders trained to classify stimulus *n-1* (green) using neuronal data sampled at two different timepoints: during stimulus *n-1* delivery (orange) or during stimulus *n* delivery (purple). Lower: validation of stimulus classifiers over a time period spanning from trial *n-1* to trial *n*. In trial *n-1*, firing rates were aligned to reward delivery while in trial *n* they were aligned to stimulus onset (vertical lines). The marginal histogram indicates stimulus frames in trial *n-1*.

How does the coordinate frame change over time? The transition could be gradual, possibly involving intermediate neuronal representations, or abrupt. To investigate this, we trained decoders on either the *n-1* (orange) or *n* (purple) epochs and evaluated their performance over time (Figure 7B, upper panel). Figure 7B (lower panel) shows decoding accuracy for *n-1*, aligned to the moment the rat crossed the light barrier in front of the spout (“reward time;” additional analyses using alternative time windows are presented in Figure S7). With this alignment, the stimulus delivery period is jittered across trials due to variability in reaction time following the go cue (inset histogram). As expected, the decoder trained on *n-1* activity (orange) performs nearly perfectly during its training window and remains accurate through reward, but its accuracy drops sharply thereafter and fails entirely during presentation of *n*. This indicates that the original representational format of *n-1* is no longer accessible.

By contrast, the decoder trained on *n-1* activity during presentation of stimulus *n* (purple) successfully recovers *n-1* in that epoch. Thus, *n-1* remains represented but in a distinct, time-shifted format. The reverse also holds: an *n*-trained decoder cannot decode *n-1* during its original presentation. Together, these results reveal two representational formats for *n-1*, each anchored to a different temporal context. One (orange) encodes *n-1* in real time, as the current sensory input; the other (purple) encodes *n-1* retrospectively, as a memory trace of the preceding episode. This suggests that the brain recruits distinct coordinate frames to represent ongoing sensory input versus recalled content, even for the same physical stimulus.

Strikingly, the transition between coordinate frames coincides with reward time. Prior to and during reward, *n-1* is treated as the present stimulus and linked to its behavioral outcome. After reward, the episode resets, and *n-1* is recoded as the previous trial’s stimulus – integrated into memory and carried forward into the next decision. This recoded trace persists through stimulus *n*, with decoding performance nearly matching that of *n* itself, and likely provides the signal that shapes the coding of *n*.

## Discussion

### Integration of Prior and Sensory Evidence in Perceptual Decisions

Perceptual decision-making requires embedding current sensory input into the recent context. We examined a task where the categorization of a stimulus is biased away from the previous stimulus (weak *n-1* leads to *n* being judged stronger, and vice versa). Bayesian inference can provide a useful description of perceptual decision-making (*1, 2*) and categorization (*59*). Applied to our experiments: sensory evidence about the current input, *n* (likelihood) combines with stored expectations (prior sensory information, approximated as *n-1*) to yield an *a posteriori* estimate of stimulus *n* category, and it is this “posterior” – rather than the raw input – that shapes percepts and behavior. We aimed to find the neuronal correlates of this convergence of present and past sensory information.

Previous work has focused on identifying the neuronal correlates of sensory priors: a representation of the sensory prior during a working-memory task has been identified in the posterior parietal cortex (*22*), a region strongly connected to vM1. A recent study (*23*) found that, in a task where mice reported whether a visual stimulus appeared on the left or right, neuronal activity during the pre-stimulus interval – across regions including primary visual cortex (V1) and the thalamic lateral geniculate nucleus – encoded information about the block-level prior. However, the presence of a neuronal representation of this prior does not establish that it contributes to the perceptual choice. The neuronal representation of the prior becomes behaviorally meaningful only when combined with current sensory evidence to influence the evolving decision. Our central question, here, is at which cortical processing stage the sensory prior (the trial-*n-1* trace) interacts with new evidence (the trial-*n* representation) to shape behavior. Our results suggest that vS1 behaves as an encoder of the likelihood of a specific stimulus category in trial *n* based on the incoming sensory input. This information is relayed by vS1 to vM1, where the likelihood is combined with a prior distribution based on previous trials, producing a posterior representation which informs the final decision.

### Single-unit vs. population coding of history effects

A neuronal population that contributes causally to the history-dependent bias should (i) encode stimulus *n*; (ii) encode stimulus *n-1*; and (iii) exhibit an *n-1*–dependent modulation of the stimulus-*n* representation whose strength and sign predict the behavioral bias. Using population decoding and trial-by-trial choice correlations, we tested vS1 and vM1 against these criteria. We included vS1 because history dependence could arise locally or be inherited from subcortical inputs, and vM1, a major frontal target of vS1 (*28–32*), as a plausible site where history-dependent modulation first aligns with choice.

Individual units in both vS1 and vM1 encoded stimulus *n* and *n-1* (i.e., firing rate covaried with vibration intensity), satisfying criteria (i) and (ii), although units meeting both simultaneously were rare. Moreover, the sign of the correlation for *n-1* did not systematically oppose that for *n* – a prerequisite for explaining the repulsive behavioral effect – indicating that individual-unit responses alone cannot account for the perceptual bias.

We therefore turned to population-level decoding, extracting information via weighted sums of unit activity. In vM1, during stimulus *n*, population activity carried information about both stimulus *n* (Figure 3A, right) and *n-1* (Figure 5A, right). vS1 robustly encoded *n* (Figure 3A, left) but carried only a marginal trace of *n-1* (Figure 5A, left). Critically, vM1 showed a systematic bias: representations of stimulus *n* were displaced by *n-1* in the repulsive direction (Figure 3C, D, right), whereas vS1 showed no such effect (Figure 3C, D, left).

Did this bias play out in behavior? To test this, we developed new choice probability measures that probe how trial-to-trial variability in neuronal activity correlates with trial-to-trial variability in decision (*47–49*). After uncovering the relationship between the decoded population state and the subsequent choice, we determined that vS1 fluctuations were not significantly predictive of choices and, most importantly, did not explain the history-dependency of choices (Figure 4E-F). In contrast, vM1 activity strongly predicted behavior: trials in which vM1’s decoding of *n* was more biased by *n-1* were more likely to yield matching repulsive perceptual reports (Figure 4F, green). Thus, only in vM1 did the influence of *n-1* on the representation of *n* align with the rats’ choices, satisfying criterion (iii). These results suggest that the repulsive *n-1* effect does not originate in vS1 or subcortical inputs but arises in downstream cortical stages, particularly vM1.

### Stages of Processing in the Transformation from Sensory Input to Contextualized Percept

Consistent with prior work (*29, 40, 60*), vS1 neurons exhibit a sharp onset response and robustly encode vibration intensity (Figure 2), reflecting short integration windows on the order of tens of milliseconds. They also display post-stimulus suppression (Figure 2), a hallmark of early sensory processing (*61*). By contrast, vM1 neuronal populations, in line with earlier reports (*37, 38, 62*), exhibit a slower, stimulus-evoked ramp, indicative of longer integration times, and lack post-stimulus suppression. These differences are consistent with a back-to-front transformation along the cortical hierarchy, characterized by progressively longer integration timescales (*63*). Nonetheless, under conditions not requiring extended integration, vM1 can respond rapidly and transiently to whisker input (*28, 64*).

Perceptual decisions may emerge through a cascade of processing stages that transform raw sensory input into contextually informed choices. Sensory receptors encode the physical properties of a stimulus with high temporal precision, without regard for functional significance or memory (*65*). In our view, early cortical stages (e.g., vS1) largely preserve this raw, temporally confined encoding (reviewed in Diamond & Toso (*66*)). Subsequent intracortical processing, we hypothesize, blends sensory input with prior experience and knowledge, thereby transforming representation of physical characteristics into perception of events embedded in the context of ongoing behavior. While vS1 carried less contextual information, our data cannot exclude vS1 involvement in memory and decision making under some conditions.

Although the classical nomenclature ties vM1 to motor output, this frontal cortical region participates in a broad range of functions beyond motor programming (*36, 39, 40, 67, 68*). Our findings identify a new role: integrating past and present sensory input to guide perceptual categorization. Recent wide-field Ca²⁺ imaging studies in mice performing tactile detection tasks (*69*) argue against the existence of cortical representations that are purely sensory (encoding only physical stimulus parameters), purely perceptual (encoding conscious sensory experience divorced from significance), or purely motor (encoding action independently of preceding events). Our results concur: vS1 provides a relatively veridical representation of stimulus intensity, only minimally influenced by contextual factors such as the previous stimuli. By contrast, vM1 encodes a representation shaped – like behavior itself – by both current sensory input *and* contextual history. vM1 may thus reflect a transitional stage linking sensory-perceptual encoding to final decision.

Though prior choices and rewards are known to affect current perceptual decisions (*10, 12–15, 17, 44, 67, 70*), earlier studies under the present study’s task conditions found only modest behavioral effects from non-stimulus history (*5*), which was factored out in our analyses (see Methods). Intriguingly, a recent study identified correlates of reward priors in vM1 (there referred to as M2) (*67*). Future work will need to examine how both stimulus and non-stimulus history converge in this brain region.

### A possible role for interneurons

One novel observation is that fast-spiking units, likely interneurons, showed opposite modulation by *n* and *n-1* (Figure 6C, Figure S6), raising the possibility that inhibitory neurons contribute directly to encoding the repulsive bias. Fast-spiking interneurons, most of which are parvalbumin-positive (PV⁺), and somatostatin-positive (SST⁺) cells – both abundant in frontal cortex (*71, 72*) – receive differential excitatory input (*73–75*) and can regulate pyramidal output in a stimulus history–dependent manner (*76–78*).

While extracellular data limit definitive cell-type identification, we speculate that joint activity across interneuron subtypes could tune the impact of *n-1* on *n* processing. A plausible mechanism involves recurrent excitation–inhibition balance within vM1, supporting slow dynamics and attractor states capable of sustaining memory traces (*79–82*).

Whether prior information is received from another brain region, or whether incoming evidence is transformed into a Bayesian posterior within the same circuit (*9, 83*), our findings support the idea that interneurons may play a distinct role in mediating the repulsive bias exerted by stimulus *n-1* on the perception of *n*, by encoding *n-1* in a manner opposite to their encoding of *n*. This is consistent with prior work showing interneurons participate in top-down modulation and gating of sensory inputs, often mediating context-dependent integration within local circuits (*53, 54, 84, 85*).

### Transition between real-time and retrospective neuronal coding

The results in Figure 7 reveal a shift in how the same stimulus, *n–1*, is represented over time in vM1. The decoder denoted by the orange trace captures *n–1* in a “real-time” frame: the activity evoked by the immediate stimulus and shortly thereafter. The purple decoder captures a “retrospective” frame: a reformatted version of *n–1* that arises between trials *n–1* and *n*. This is not a decaying echo – its decoding accuracy remains steady during trial *n*; memory and perception are both coded robustly. These two formats are not interchangeable: a decoder trained on *n–1* activity cannot recover that stimulus during *n*, and vice versa. This suggests that the brain does not simply preserve or degrade a static code but shifts the coordinate system itself.

This shift best aligns with reward delivery. Before reward, *n–1* is encoded as the choice-relevant input linked to the outcome. After reward, the same stimulus appears as a memory trace. The representational transition around reward time suggests that the brain recodes recent stimuli into a new coordinate frame, transforming “just-experienced input” into “remembered content.” As such, *n–1* enters trial *n* not in its original form, but recast in the context of a new episode. This memory trace shapes the representation of *n* and influences the decision that follows (Figure 5).

The segmentation and representational shift observed in vM1 parallels mechanisms long associated with the hippocampus and episodic memory (*86*). Hippocampal output may help initiate or stabilize the coordinate frame transition we observe in vM1 – especially given established loops between hippocampus and frontal/motor areas (*87*). Alternatively, vM1 may perform this transformation autonomously, with episodic structure emerging locally to serve the demands of decision-making.

## Materials and Methods

### Experimental Design

#### Subjects

Subjects were male Wistar rats (Harlan Laboratories, San Pietro Al Natisone). For the neurophysiological results, data from five rats were obtained, while four additional rats provided only behavioral data (Figure S1). All rats were handled and trained daily. Initially, they were housed in pairs—either with each other or with rats not participating in the present experiments. Prior to microdrive implantation (see below), they were moved to individual housing. Rats were regularly monitored for health and welfare, and provided with daily environmental and social enrichment. They were maintained on a 12/12 h light/dark cycle. To promote motivation during the behavioral task, rats were water-restricted between daily testing sessions, while having continuous free access to food. On days without experimental sessions, water was freely available. Testing occurred on each working day in ∼1 h sessions. Water was provided during behavioral sessions in the form of a sweet fluid reward. After each session, rats were offered free access to plain water to ensure satiation.

#### Behavioral task

An LED positioned above the nose poke signaled trial availability. A trial began when the rat crossed the optical sensor within the nose poke, in this position, the rat’s right whiskers touched the plate (Figure 1A). After nose poke activation, a 400 ms delay preceded a 500 ms plate vibration. Following this vibration, a variable delay (400–600 ms) occurred before an auditory “go” cue instructed the rat to withdraw and choose one of two reward spouts. Choice was detected by infrared beams at the entrance to each spout. The rewarded side was determined by vibration intensity (see “Vibrotactile stimuli”): e.g., intensities above the category boundary (“strong”) rewarded the left (right) spout, while intensities below (“weak”) rewarded the right (left) spout. Incorrect choices received no reward. For trials at the category boundary, reward delivery was randomized between the two spouts. After an incorrect choice, the nose poke sensor was disabled for 1500–3500 ms, enforcing a short waiting period before the next trial. To discourage development of side biases, after three consecutive intensities rewarded on one side, the fourth trial was always sampled from the set of intensities rewarded on the opposite side. This led to a slightly above-chance probability of alternation between subsequent stimulus categories (56.8%). Training or testing sessions typically lasted ∼1 h and contained ∼300 trials. When a rat achieved >75% correct performance for five consecutive sessions, training was considered complete and subsequent sessions were used for analysis.

#### Vibrotactile stimuli

Stimuli were delivered via a rectangular plate (20 × 30 mm) mounted on a motorized shaker (Bruel & Kjaer, type 4808), driven by analog velocity signals to produce movement along the rostro-caudal axis. Vibrations consisted of low-pass–filtered white noise (as in Hachen et al.(*5*)). For each trial, velocity values were sampled at 10 kHz from a zero-mean normal distribution. The noise signal was low-pass filtered (Butterworth, 150 Hz cutoff), amplified, and sent as voltage input to the shaker motor. Vibration intensity (“mean speed”) was quantified as the nominal mean absolute velocity, calculated as the standard deviation of the velocity distribution multiplied by √(2/π). This measure has been identified as supporting high-acuity discrimination of whisker-mediated vibrations in rats (*88*). In each session, nine linearly spaced intensity values were used. On each trial the software randomly selected one of these nine intensities with a probability of 0.11. For clarity, intensity was converted from mean speed (in mm/s) to a scale of −4 (lowest intensity) to 4 (highest intensity), with the reward rule boundary at 0.

For each velocity standard deviation (intensity), 50 unique “seeds” (time series) were available. Because each vibration’s velocity profile was generated by random sampling at 10 kHz from a normal distribution, no two seeds were identical. Thus, even stimuli with the same nominal intensity differed in actual velocity profile.

### Experimental apparatus

The apparatus (custom-built by CyNexo, https://www.cynexo.com/) consisted of a Plexiglas box (25 × 25 × 38 cm, height × width × length) housed within a sound- and light-attenuated chamber. A rounded head-port on the front wall provided access to the circular nose poke aperture (0.85 cm diameter). On each side of the box, a metal reward spout with a plastic lip delivered 0.03 mL of water-diluted juice via syringe pumps upon correct choices. During sessions, the box was closed with a Plexiglas cover and monitored via an overhead camera. Task control software for both rat and human experiments was written in-house in LabVIEW (National Instruments, Austin, TX).

#### Implantation of microdrives and neuronal recordings

Custom multi-electrode microdrive arrays were implanted for chronic electrophysiological recordings. The arrays (aoDrive, CyNexo; https://www.cynexo.com/portfolio/neural-drives/) incorporated either 11 or 15 single tungsten electrodes (FHC) with 400 μm spacing, plus an optic fiber. The microdrives and TDT connectors were fixed to the skull using dental cement. Electrophysiological recordings began 7–10 days after surgery, once animals had fully recovered, during which they had *ad libitum* access to food and water and were monitored daily. Recording quality was maintained for 3–6 months post-implantation.

Recordings were obtained from five rats, targeting vS1 and/or vM1. A TDT system (Tucker-Davis Technologies) was used for pre-amplification, filtering, and digitization of extracellular signals at 24 kHz. Position sensor signals were also recorded for synchronization with behavioral events. At the end of each recording session, electrodes were advanced or retracted by 50 μm to sample new neuronal populations across cortical layers. In ∼40% of sessions, vS1 and vM1 were recorded simultaneously; in the remainder, only one area was recorded.

### Quantification and statistical analysis

#### Analysis of behavioral data

Data analysis was performed in MATLAB 2022b (MathWorks, Natick MA). For each stimulus intensity, we counted the number of times that the rat responded by turning to the side corresponding to “weak” or “strong”, then we fitted the responses with psychometric curves. For the analyses related to Figure 1B-C and Figure S1, as in Hachen et al. (*5*), data was fitted with a cumulative Gaussian function with lapse parameters (*89*):

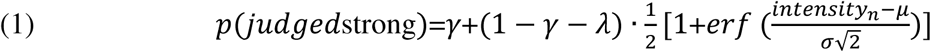

Where *μ* is the midpoint parameter, *σ* the slope (standard deviation) parameter, *γ* and *λ* the lower and higher lapse parameters. Parameter values were estimated by maximum likelihood using the MATLAB function fmincon.

When modeling the impact of both intensity *n* and *n-1* on choice *n* (Figure 1D), the argument of the cumulative Gaussian function was modified to include intensity in both trial *n* and in trial *n-1* as linear predictors:

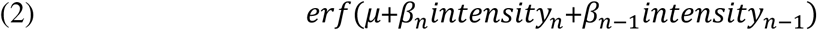

Where *μ* is the midpoint parameter, *β*_*n*_ a linear coefficient representing the influence of the intensity received in trial *n, β*_*n*−1_ a linear coefficient representing the influence of intensity *n-1*.

The variance of parameter estimates and model predictions was computed by means of parametric bootstrapping: after calculating the probability of judging each stimulus *n* as “strong” by fitting the model, we simulated binary decisions drawing from this probability distribution. This procedure was repeated 100 times. For each of these 100 repetitions the psychometric model was re-fitted to the new sample of simulated binary decisions, producing a slightly different output every time. The 95% confidence intervals on model predictions are shown in Figure 1D (light shading).

To exclude that the observed repulsive effect of intensity *n-1* on choice *n* could result from above-chance alternation between the intensity categories of stimuli *n-1* and *n*, we performed the following control analysis. First, we fitted the data with a simplified model including only intensity *n* as a term (i.e. without intensity *n-1*), and generated artificial choices by binarizing the simplified model’s predictions. We generated as many choices as in the original dataset, preserving the exact same trial sequence, and repeated the procedure 100 times. If the mere sequence of stimuli were responsible for the measured effect of intensity *n-1*, then an effect of *n-1* should be observed also in this artificial dataset. We therefore fitted the artificial dataset with the original model with predictors for both intensity *n* and *n-1*, but the resulting coefficient for *n-1* was null (Figure S1B, gray line).

#### Neuronal data processing

Spike sorting was performed using Kilosort 2.0 (*90*) on raw voltage traces. Resulting clusters were manually curated in Phy 2.0 (*91*) and classified as single-unit or multi-unit based on waveform shape, autocorrelogram, and the similarity index from Kilosort. From vS1, we obtained 264 units from 28 sessions in 5 rats with a mean of 9 ± 6 (s.d.) units/session. From vM1, we obtained 594 units from 35 sessions in 3 rats (17 ± 7 units/session). This difference partly reflects variability in the number of effective recording channels, influenced by cranial anatomy.

Spike trains were aligned to behavioral events using position sensors and stimulus timing signals. For each sorted unit, mean waveforms were computed, and spike width was measured as the interval between the negative trough and the subsequent positive peak (*56–58*). Units with widths <0.4 ms were classified as putative fast-spiking neurons; those >0.6 ms as putative regular-spiking neurons. We did not label neurons in which the measured spike width exceeded 1 ms, and neurons with an uncommonly high standard deviation in their waveform (>95^th^ percentile), likely to be corrupted by noise artifacts. All processing was performed in custom MATLAB (2022) and Python 3 scripts.

#### Analysis of stimulus tuning in individual units

Analyses were performed in MATLAB 2022 (MathWorks). Tuning to stimulus *n* and stimulus *n – 1* was quantified by Spearman’s rank correlation between the normalized nominal intensity (–4 to 4, step 1) and firing rate in a 500 ms window aligned to stimulus presentation.

To remove collinearity between stimulus *n – 1* and choice *n – 1*, correlations were computed separately for trials conditioned on the choice in trial *n – 1* (“weak” or “strong”). This procedure resulted in two separate coefficients for stimulus *n-1* and, in analogy, two separate coefficients for stimulus *n*, which were then averaged. Significance was assessed via a permutation test: the same analysis was repeated after 1000 random permutations of the intensity labels, generating a null distribution. Units were considered significantly tuned if their averaged correlation coefficient lay outside the 95% two-tailed confidence interval of this null distribution.

#### Decoding of task variables from neuronal populations

We decoded trial conditions from population firing rates using a pseudo-simultaneous population approach (*92, 93*). This method was chosen because the number of simultaneously recorded units varied (2 to 32) and recordings spanned multiple laminar depths. Combining units recorded separately allowed us to capture contributions from diverse neuronal types and account for sparse coding across the population. Additionally, resampling ensured balanced trial conditions within behavioral sessions.

Only units with an average firing rate above 1 Hz (∼85% of units) were included. Where stated in the text (e.g. for the analyses shown in Figure 3), units were pre-selected based on significant tuning to stimulus *n* (Spearman’s correlation; see Figure 2c). In all other analyses, all units meeting the basic criteria were used regardless of tuning. Decoding combined units across rats after verifying consistent results at the individual rat level when sample size allowed it.

Decoding was performed separately for each brain area (vS1 or vM1), as follows:

1. **Labeling trials:** For each unit, trials were labeled by the condition of interest. For stimulus *n*, trials were categorized as “strong” or “weak” vibration, excluding ∼11% of trials with stimulus on the boundary. Trials were also labeled by stimulus *n–1* category, forming a 2×2 design matrix (Figure S3A). The same procedure was adopted when decoding choice *n*, employing “strong/weak” decision as a label instead of stimulus *n*. When decoding stimulus *n-1*, the 2 factors of the 2×2 design matrix were stimulus *n-1* (“strong/weak”) and choice *n-1* (“strong/weak”), thus balancing choice *n-1* to control for any stimulus-unrelated serial effects (see main text). Matrices with more than 2 factors (e.g. including both trials *n-1* and *n-2*) were infeasible due to trial count limits. If the experimental session contained more than 300 trials, only the first 300 trials were analyzed to ensure high animal motivation. For decoding, 70% of trials were used for training and 30% (out-of-training sample) for validation, balanced across conditions. Trial counts per neuron ranged from 18–75 (5–25 validation trials) for stimulus and choice *n*, fewer for stimulus *n–1* due to error trial inclusion (∼20–25%), but never below 9 trials per condition. Rats had opposite response contingencies (“left” or “right” mapped to “weak” or “strong”), so units were further sampled to balance left/right responses, minimizing potential confounds from movement- or side-related activity.
2. **Firing rate resampling:** For each selected unit and training trial, firing rates were computed in a 400 ms time window. Rates were resampled with replacement 4000 times, drawing 1000 trials from each of the 4 conditions of the relevant design matrix. This procedure created vectors ***f*** of firing rates sorted by condition. Another vector ***f*** was computed analogously by resampling validation trials. The procedure was repeated for all the units selected in step 1).
3. **Pseudo-trial construction:** Resampled firing rate vectors from all units were concatenated into matrices (***f***-by-***u***, where ***f***=4000 resamples and ***u*** is the total number of units included in the analysis; Figure S3A). This procedure was performed separately with the training and the validation vectors, yielding two different ***f***-by-***u*** matrices. Firing rates were *z*-scored within each unit.
4. **Classification:** A linear binary Support Vector Machine (SVM) (*94*) was trained on the ***f***-by-***u*** training matrix, with each row *f*_*i*_ considered as an observation in *u* dimensions corresponding to a pseudo-simultaneous population state, yielding 2000 observations per class. The second factor of the design matrix (e.g. stimulus *n-1*) was balanced and hidden to the classifier. The SVM identified the optimal separating hyperplane:

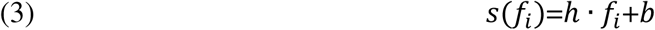 where ***h*** is the ***u***-dimensional vector of hyperplane coefficients, *f_i_* an observation in population space, and ***b*** a bias term. Observations with *s*(*f_i_*)>0 were classified as the positive class (e.g., “strong” stimulus), and those with *s*(*f_i_*)>0 as negative (“weak”). The *s*(*f_i_*) represents the distance from the hyperplane (the “score”). The trained hyperplane was then applied to the validation set. To assess the contribution of each unit in defining the orientation of the hyperplane, we considered the vector *w*, orthogonal to the hyperplane’s coefficient vector *h*, containing a scalar weight for each unit *u_j_* in the population. This decoding was repeated across 400 ms windows shifted in 25 ms steps, spanning –400 to 900 ms relative to stimulus onset (–400 to 0 ms pre-stimulus; 500 to 900 ms post-stimulus). For each iteration, new training and validation samples were drawn. We cross-validated 100 decoders and averaged accuracy, increasing to 500 decoders for Figure 4 to better estimate variability. As control, labels were shuffled and decoding repeated to assess significance by comparing accuracies from shuffled versus non-shuffled data, estimating the probability that observed accuracy exceeded chance. This measure estimates the probability that a randomly sampled accuracy value resulting from non-shuffled labels was higher than a randomly sampled accuracy value resulting from shuffled labels, testing all possible combinations. To verify no confounds from trial transitions, we repeated decoding swapping trial *n–1* with trial *n+1*, which should be unpredictable and irrelevant; accordingly, decoding accuracy dropped to chance-level.

### Marginalization of task variables within population states

As described previously (step 3), decoding used an *f*-by-*u* matrix of pseudo-simultaneous population states (Figure S3A), where *f* is the number of resampled firing rate samples and *u* the number of units. For each firing rate matrix, we created a corresponding *f*-by-*u* matrix encoding the rat’s choice in each resampled trial (Figure S3B), with choices coded as 0 (“weak”) or 1 (“strong”), analogous to psychometric analyses.

Each row *f_i_* of the decoding matrix represents a population state, and the corresponding row in the choice matrix represents the distribution of behavioral choices given that state. Marginalizing across units (averaging over the *u* columns) yields the expected probability the rats report stimulus *n* as “strong” for that population state:

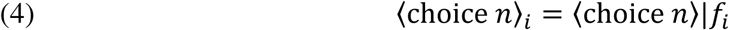

where 〈choice *n*〉 refers to the expected value of choice *n* across all units *u*.

Similarly, other task variables (e.g., stimulus *n* and *n–1* values) were stored in analogous *f*-by-*u* matrices and marginalized over units to estimate their expected values per population state:

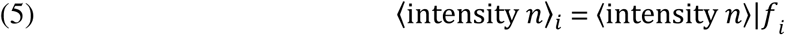

where 〈intensity *n*〉 refers to the expected value of intensity *n* across all units *u*.

We refer to the vectors storing all the values of marginalized task variables as ⟨*variable*⟩. For each value of the marginalized variable, ⟨*variable*⟩_*i*_, there is a corresponding *score_i_* resulting from population decoding (Figure S3A-B).

In Fig. 4C-D, to control for the influence of the intensity value and the stimulus seed on the rat’s choice, we selected only pseudo-simultaneous trials in which the marginalized *n* and *n-1* stimulus values were within a range of (-)2.4 and (-)2.5 intensity.

### Linear regression of choice with pseudo-simultaneous population states

To assess how population readouts relate to the rat’s choices, we considered each of the cross-validated decoders (500 in Fig. 4A, 100 in Fig. 5C), each tested on 4,000 pseudo trials (totaling 400,000 population states). For each decoder, two linear regression models were fitted to predict the frequency of “strong” choices from each decoded trial. Both models included marginalized intensity values of stimulus *n* and *n–1* as predictors. Beyond stimulus *n* and *n–1*, the “population-informed” model also included the decoder’s output score for each population state (higher scores indicate stronger decoder confidence in classifying the stimulus as “strong”):

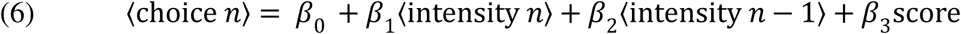

where each term is weighted with a linear coefficient *β*; *β*_o_ is an intercept term. The score refers to the decoded stimulus category in either *n* or *n–1*, depending on the analysis. By contrast, the population-blind model omitted the score term:

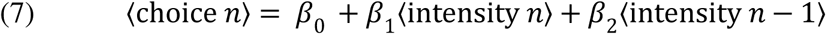

Models were fitted separately for each 400 ms time window used in decoding. For each population state *f_i_*, the population-blind model yielded the same prediction regardless of decoding time, due to the absence of the score term.

Model efficiency was compared via Akaike Information Criterion (AIC), as it balances goodness of fit with model complexity (two parameters for “uninformed” but three parameters for “informed”). Between two models, the one with the lower AIC is more effective at explaining the observed data, even after accounting for model complexity (*50*). Because models were fitted to resampled data, AIC differences were corrected through permutation testing: informed and uninformed models were fit to data with shuffled trial labels, and resulting AIC differences were subtracted from original differences. Finally, these permutation-corrected AIC differences were averaged across decoders to yield the time series shown in Figs. 4A and 5C.

### Quantification of shared variability in history bias between neuronal decoding and behavior

To quantify shared variability in history-dependent bias (Fig. 4F), we analyzed 500 decoders with distinct cross-validation samples. Each decoder’s validation set (30% of trials) introduced variability in decoding and observed behavioral bias. We measured decoder bias as the difference in average output score between trials where stimulus *n–1* was “strong” versus “weak.” Behavioral bias was the difference in marginal choice probability under the same conditions. This yielded two 500-element vectors (decoder bias and behavioral bias), one per decoder. Their relationship was quantified across decoders by Pearson’s correlation coefficient, repeated across all decoding time windows, and its significance tested with a permutation test (see Results).

## Acknowledgments

We thank several members of the research team for valuable discussions, including Arash Fassihi, Nader Nikbakht, Francesca Pulecchi, Jacopo Rigosa, and Zahra Yousefi Darani. We thank Marco Gigante (SISSA Mechatronics) and Fabrizio Manzino (CyNexo) for their invaluable technical assistance. Mauro Dall’Argine, Mario Fontanini and Sara Mohammed assisted in animal training and behavioral data acquisition. We are grateful for valuable feedback from Iacopo Hachen’s PhD thesis committee and from Tommaso Fellin. We are particularly grateful to Stefano Panzeri for supervising Alejandro Pequeno-Zurro’s PhD thesis and providing inputs on data curation and preliminary analyses of tactile neuronal coding.

## Funding

Human Frontier Science Program, project RGP0017/2021 (MD)

Italian Ministry PRIN 2022 contract 20224FWF2J (MD)

Italian Ministry PNRR contract P20229752W (MD)

European Union MSCA postdoctoral fellowship grant 101153670 (IH)

NARSAD Young Investigator Grant from the Brain & Behavior Research Foundation 30389 (SR)

Regional Laboratory for Advanced Mechatronics, LAMA FVG (http://lamafvg.it)

European Union Horizon 2020 MSCA Programme grant 813713 (MED, APZ)

Ubbo Emmius Funds of the University of Groningen (APZ)

## Author contributions

Conceptualization: IH, SR, & MED

Data curation: AS, APZ, IH, SR

Formal analysis: IH

Funding acquisition: MED

Investigation: SR, IH, AS

Methodology: SR, IH

Project administration: MED, IH

Resources: MED, SR

Software: IH, AS, APZ

Supervision: MED

Validation: IH, SR, AS, APZ

Visualization: IH

Writing – original draft: IH & MED

Writing – review & editing: IH, SR, & MED

## Competing interests

Authors declare that they have no competing interests.

## Data and materials availability

Data will be released on an Open Science Framework repository upon publication of the present manuscript in a peer-reviewed journal. The datasets will be provided in standardized formats, with metadata and usage documentation. Codes with usage documentation will be released under an open license upon publication of the present manuscript in a peer-reviewed journal.

## Notes

### Competing Interest Statement

The authors have declared no competing interest.

### Summary of Updates

- Updated/clarified acknowledgments. - Removed a duplicate in the references.

## References

1. A. Pouget, J. M. Beck, W. J. Ma, P. E. Latham, Probabilistic brains: knowns and unknowns. Nature Neuroscience 16, 1170–1178 (2013).

2. W. J. Ma, M. Jazayeri, Neural Coding of Uncertainty and Probability. Annual Review of Neuroscience 37, 205–220 (2014).

3. W. J. Ma, Bayesian Decision Models: A Primer. Neuron 104, 164–175 (2019).

4. A. Fassihi, A. Akrami, V. Esmaeili, M. E. Diamond, Tactile perception and working memory in rats and humans. PNAS 111, 2331–2336 (2014).

5. I. Hachen, S. Reinartz, R. Brasselet, A. Stroligo, M. E. Diamond, Dynamics of history-dependent perceptual judgment. Nat Commun 12, 6036 (2021).

6. C. Waiblinger, C. M. Wu, M. F. Bolus, P. Y. Borden, G. B. Stanley, Stimulus Context and Reward Contingency Induce Behavioral Adaptation in a Rodent Tactile Detection Task. J. Neurosci. 39, 1088–1099 (2019).

7. P. Ashourian, Y. Loewenstein, Bayesian Inference Underlies the Contraction Bias in Delayed Comparison Tasks. PLoS One 6 (2011).

8. O. Raviv, I. Lieder, Y. Loewenstein, M. Ahissar, Contradictory Behavioral Biases Result from the Influence of Past Stimuli on Perception. PLOS Computational Biology 10, e1003948 (2014).

9. H. Sohn, D. Narain, N. Meirhaeghe, M. Jazayeri, Bayesian Computation through Cortical Latent Dynamics. Neuron 103, 934–947.e5 (2019).

10. M. Lages, M. Treisman, Spatial frequency discrimination: Visual long-term memory or criterion setting? Vision Research 38, 557–572 (1998).

11. S. Gepshtein, M. Kubovy, Stability and change in perception: spatial organization in temporal context. Exp Brain Res 160, 487–495 (2005).

12. A. E. Urai, J. W. de Gee, K. Tsetsos, T. H. Donner, Choice history biases subsequent evidence accumulation. eLife 8, e46331 (2019).

13. A. Braun, A. E. Urai, T. H. Donner, Adaptive History Biases Result from Confidence-Weighted Accumulation of past Choices. J. Neurosci. 38, 2418–2429 (2018).

14. A. Lak, E. Hueske, J. Hirokawa, P. Masset, T. Ott, A. E. Urai, T. H. Donner, M. Carandini, S. Tonegawa, N. Uchida, A. Kepecs, Reinforcement biases subsequent perceptual decisions when confidence is low, a widespread behavioral phenomenon. eLife 9, e49834 (2020).

15. L. Busse, A. Ayaz, N. T. Dhruv, S. Katzner, A. B. Saleem, M. L. Schölvinck, A. D. Zaharia, M. Carandini, The Detection of Visual Contrast in the Behaving Mouse. J. Neurosci. 31, 11351–11361 (2011).

16. K. Iigaya, M. S. Fonseca, M. Murakami, Z. F. Mainen, P. Dayan, An effect of serotonergic stimulation on learning rates for rewards apparent after long intertrial intervals. Nature Communications 9, 2477 (2018).

17. A. Hermoso-Mendizabal, A. Hyafil, P. E. Rueda-Orozco, S. Jaramillo, D. Robbe, J. de la Rocha, Response outcomes gate the impact of expectations on perceptual decisions. Nature Communications 11, 1057 (2020).

18. R. Nogueira, J. M. Abolafia, J. Drugowitsch, E. Balaguer-Ballester, M. V. Sanchez-Vives, R. Moreno-Bote, Lateral orbitofrontal cortex anticipates choices and integrates prior with current information. Nat Commun 8, 14823 (2017).

19. V. Rao, G. C. DeAngelis, L. H. Snyder, Neural Correlates of Prior Expectations of Motion in the Lateral Intraparietal and Middle Temporal Areas. J. Neurosci. 32, 10063–10074 (2012).

20. Z. Yousefi-Darani, I. Hachen, M. E. Diamond, Dynamics of the judgment of tactile stimulus intensity. Neuromorph. Comput. Eng. 3, 014014 (2023).

21. F. Schönsberg, D. Giana, Y. Chopra, M. E. Diamond, S. Goldt, Diverse perceptual biases emerge from Hebbian plasticity in a recurrent neural network model. Neuron 113, 3673–3684.e6 (2025).

22. A. Akrami, C. D. Kopec, M. E. Diamond, C. D. Brody, Posterior parietal cortex represents sensory history and mediates its effects on behaviour. Nature 554, 368–372 (2018).

23. C. Findling, F. Hubert, L. Acerbi, B. Benson, J. Benson, D. Birman, N. Bonacchi, E. K. Buchanan, S. Bruijns, M. Carandini, J. A. Catarino, G. A. Chapuis, A. K. Churchland, Y. Dan, F. Davatolhagh, E. E. J. DeWitt, T. A. Engel, M. Fabbri, M. A. Faulkner, I. R. Fiete, L. Freitas-Silva, B. Gerçek, K. D. Harris, M. Häusser, S. B. Hofer, F. Hu, J. M. Huntenburg, A. Khanal, C. Krasniak, C. Langdon, C. A. Langfield, P. E. Latham, P. Y. P. Lau, Z. Mainen, G. T. Meijer, N. J. Miska, T. D. Mrsic-Flogel, J.-P. Noel, K. Nylund, A. Pan-Vazquez, L. Paninski, J. Pillow, C. Rossant, N. Roth, R. Schaeffer, M. Schartner, Y. Shi, K. Z. Socha, N. A. Steinmetz, K. Svoboda, C. Tessereau, A. E. Urai, M. J. Wells, S. J. West, M. R. Whiteway, O. Winter, I. B. Witten, A. Zador, Y. Zhang, P. Dayan, A. Pouget, Brain-wide representations of prior information in mouse decision-making. Nature 645, 192–200 (2025).

24. P. Kok, J. F. M. Jehee, F. P. de Lange, Less Is More: Expectation Sharpens Representations in the Primary Visual Cortex. Neuron 75, 265–270 (2012).

25. A. G. Bondy, R. M. Haefner, B. G. Cumming, Feedback determines the structure of correlated variability in primary visual cortex. Nat Neurosci 21, 598–606 (2018).

26. J. H. Macke, H. Nienborg, Choice (-history) correlations in sensory cortex: cause or consequence? Current Opinion in Neurobiology 58, 148–154 (2019).

27. M. Fritsche, S. G. Solomon, F. P. de Lange, Brief Stimuli Cast a Persistent Long-Term Trace in Visual Cortex. J. Neurosci. 42, 1999–2010 (2022).

28. J. Smith, K. Alloway, Rat whisker motor cortex is subdivided into sensory-input and motor-output areas. Frontiers in Neural Circuits 7 (2013).

29. V. Esmaeili, K. Tamura, G. Foustoukos, A. Oryshchuk, S. Crochet, C. C. Petersen, Cortical circuits for transforming whisker sensation into goal-directed licking. Current Opinion in Neurobiology 65, 38–48 (2020).

30. C. C. H. Petersen, Cortical Control of Whisker Movement. Annual Review of Neuroscience 37, 183–203 (2014).

31. I. Ferezou, F. Haiss, L. J. Gentet, R. Aronoff, B. Weber, C. C. H. Petersen, Spatiotemporal Dynamics of Cortical Sensorimotor Integration in Behaving Mice. Neuron 56, 907–923 (2007).

32. F. Matyas, V. Sreenivasan, F. Marbach, C. Wacongne, B. Barsy, C. Mateo, R. Aronoff, C. C. H. Petersen, Motor Control by Sensory Cortex. Science 330, 1240–1243 (2010).

33. M. Brecht, M. Schneider, B. Sakmann, T. W. Margrie, Whisker movements evoked by stimulation of single pyramidal cells in rat motor cortex. Nature 427, 704–710 (2004).

34. C. L. Ebbesen, G. Doron, C. Lenschow, M. Brecht, Vibrissa motor cortex activity suppresses contralateral whisking behavior. Nat Neurosci 20, 82–89 (2017).

35. M. Brecht, Movement, Confusion, and Orienting in Frontal Cortices. Neuron 72, 193–196 (2011).

36. C. L. Ebbesen, M. N. Insanally, C. D. Kopec, M. Murakami, A. Saiki, J. C. Erlich, More than Just a “Motor”: Recent Surprises from the Frontal Cortex. J. Neurosci. 38, 9402–9413 (2018).

37. J. C. Erlich, M. Bialek, C. D. Brody, A Cortical Substrate for Memory-Guided Orienting in the Rat. Neuron 72, 330–343 (2011).

38. M. Murakami, H. Shteingart, Y. Loewenstein, Z. F. Mainen, Distinct Sources of Deterministic and Stochastic Components of Action Timing Decisions in Rodent Frontal Cortex. Neuron 94, 908–919.e7 (2017).

39. T. D. Hanks, C. D. Kopec, B. W. Brunton, C. A. Duan, J. C. Erlich, C. D. Brody, Distinct relationships of parietal and prefrontal cortices to evidence accumulation. Nature 520, 220–223 (2015).

40. A. Fassihi, A. Akrami, F. Pulecchi, V. Schönfelder, M. E. Diamond, Transformation of Perception from Sensory to Motor Cortex. Current Biology 27, 1585–1596.e6 (2017).

41. F. Barthas, A. C. Kwan, Secondary Motor Cortex: Where ‘Sensory’ Meets ‘Motor’ in the Rodent Frontal Cortex. Trends in Neurosciences 40, 181–193 (2017).

42. R. Quian Quiroga, S. Panzeri, Extracting information from neuronal populations: information theory and decoding approaches. Nat Rev Neurosci 10, 173–185 (2009).

43. M. Rigotti, O. Barak, M. R. Warden, X.-J. Wang, N. D. Daw, E. K. Miller, S. Fusi, The importance of mixed selectivity in complex cognitive tasks. Nature 497, 585–590 (2013).

44. M. Fritsche, P. Mostert, F. P. de Lange, Opposite Effects of Recent History on Perception and Decision. Current Biology 27, 590–595 (2017).

45. H. Safaai, M. von Heimendahl, J. M. Sorando, M. E. Diamond, M. Maravall, Coordinated Population Activity Underlying Texture Discrimination in Rat Barrel Cortex. J. Neurosci. 33, 5843–5855 (2013).

46. D. H. O’Connor, S. P. Peron, D. Huber, K. Svoboda, Neural Activity in Barrel Cortex Underlying Vibrissa-Based Object Localization in Mice. Neuron 67, 1048–1061 (2010).

47. Y. Zuo, H. Safaai, G. Notaro, A. Mazzoni, S. Panzeri, M. E. Diamond, Complementary Contributions of Spike Timing and Spike Rate to Perceptual Decisions in Rat S1 and S2 Cortex. Current Biology 25, 357–363 (2015).

48. S. Panzeri, C. D. Harvey, E. Piasini, P. E. Latham, T. Fellin, Cracking the Neural Code for Sensory Perception by Combining Statistics, Intervention, and Behavior. Neuron 93, 491–507 (2017).

49. G. Pica, E. Piasini, H. Safaai, C. A. Runyan, M. E. Diamond, T. Fellin, C. Kayser, C. D. Harvey, S. Panzeri, Quantifying how much sensory information in a neural code is relevant for behavior. Advances in Neural Information Processing Systems, 3689–3699 (2017).

50. K. P. Burnham, D. R. Anderson, Eds., Model Selection and Multimodel Inference (Springer, New York, NY, ed. 2nd, 2002; http://link.springer.com/10.1007/b97636).

51. V. de Lafuente, R. Romo, Neuronal correlates of subjective sensory experience. Nat Neurosci 8, 1698–1703 (2005).

52. R. Hattori, K. V. Kuchibhotla, R. C. Froemke, T. Komiyama, Functions and dysfunctions of neocortical inhibitory neuron subtypes. Nat Neurosci 20, 1199–1208 (2017).

53. K. V. Kuchibhotla, J. V. Gill, G. W. Lindsay, E. S. Papadoyannis, R. E. Field, T. A. H. Sten, K. D. Miller, R. C. Froemke, Parallel processing by cortical inhibition enables context-dependent behavior. Nat Neurosci 20, 62–71 (2017).

54. K. Kuchibhotla, B. Bathellier, Neural encoding of sensory and behavioral complexity in the auditory cortex. Current Opinion in Neurobiology 52, 65–71 (2018).

55. F. Studer, T. R. Barkat, Inhibition in the auditory cortex. Neuroscience & Biobehavioral Reviews 132, 61–75 (2022).

56. A. Lak, E. Arabzadeh, M. E. Diamond, Enhanced Response of Neurons in Rat Somatosensory Cortex to Stimuli Containing Temporal Noise. Cerebral Cortex 18, 1085–1093 (2008).

57. Z. V. Guo, N. Li, D. Huber, E. Ophir, D. Gutnisky, J. T. Ting, G. Feng, K. Svoboda, Flow of Cortical Activity Underlying a Tactile Decision in Mice. Neuron 81, 179–194 (2014).

58. M. Solyga, T. R. Barkat, Emergence and function of cortical offset responses in sound termination detection. eLife 10, e72240 (2021).

59. H. Lee, H.-J. Lee, K. W. Choe, S.-H. Lee, Neural Evidence for Boundary Updating as the Source of the Repulsive Bias in Classification. J. Neurosci. 43, 4664–4683 (2023).

60. S. Reinartz, A. Fassihi, M. Ravera, L. Paz, F. Pulecchi, M. Gigante, M. E. Diamond, Direct contribution of the sensory cortex to the judgment of stimulus duration. Nat Commun 15, 1712 (2024).

61. M. Maravall, “Adaptation and Sensory Coding” in Principles of Neural Coding (CRC Press, Boca Raton, FL, 2013), pp. 357–377.

62. C. D. Kopec, J. C. Erlich, B. W. Brunton, K. Deisseroth, C. D. Brody, Cortical and Subcortical Contributions to Short-Term Memory for Orienting Movements. Neuron 88, 367–377 (2015).

63. R. Rossi-Pool, A. Zainos, M. Alvarez, S. Parra, J. Zizumbo, R. Romo, Invariant timescale hierarchy across the cortical somatosensory network. Proceedings of the National Academy of Sciences 118, e2021843118 (2021).

64. T. Yamashita, A. Pala, L. Pedrido, Y. Kremer, E. Welker, C. C. H. Petersen, Membrane Potential Dynamics of Neocortical Projection Neurons Driving Target-Specific Signals. Neuron 80, 1477–1490 (2013).

65. E. Arabzadeh, E. Zorzin, M. E. Diamond, Neuronal Encoding of Texture in the Whisker Sensory Pathway. PLoS Biology 3, e17–e17 (2005).

66. M. E. Diamond, A. Toso, Tactile cognition in rodents. Neuroscience & Biobehavioral Reviews 149, 105161 (2023).

67. K. Ishizu, S. Nishimoto, Y. Ueoka, A. Funamizu, Localized and global representation of prior value, sensory evidence, and choice in male mouse cerebral cortex. Nat Commun 15, 4071 (2024).

68. D. Gupta, C. D. Kopec, A. G. Bondy, T. Z. Luo, V. A. Elliott, C. D. Brody, A multi-region recurrent circuit for evidence accumulation in rats. bioRxiv [Preprint] (2024). 10.1101/2024.07.08.602544.

69. M. Sokoletsky, A. A. Nikitina, K. Waychal, Y. Katz, I. Lampl, Inherent coupling of perceptual judgments to actions in the mouse cortex. bioRxiv, 2025.10.30.685660 (2025).

70. D. Pascucci, G. Mancuso, E. Santandrea, C. Della Libera, G. Plomp, L. Chelazzi, Laws of concatenated perception: Vision goes for novelty, decisions for perseverance. PLoS Biol 17 (2019).

71. B. Rudy, G. Fishell, S. Lee, J. Hjerling-Leffler, Three groups of interneurons account for nearly 100% of neocortical GABAergic neurons. Developmental Neurobiology 71, 45–61 (2011).

72. R. Tremblay, S. Lee, B. Rudy, GABAergic Interneurons in the Neocortex: From Cellular Properties to Circuits. Neuron 91, 260–292 (2016).

73. A. J. Apicella, I. R. Wickersham, H. S. Seung, G. M. G. Shepherd, Laminarly Orthogonal Excitation of Fast-Spiking and Low-Threshold-Spiking Interneurons in Mouse Motor Cortex. J. Neurosci. 32, 7021–7033 (2012).

74. X. Xu, E. M. Callaway, Laminar Specificity of Functional Input to Distinct Types of Inhibitory Cortical Neurons. J. Neurosci. 29, 70–85 (2009).

75. S. J. Cruikshank, T. J. Lewis, B. W. Connors, Synaptic basis for intense thalamocortical activation of feedforward inhibitory cells in neocortex. Nat Neurosci 10, 462–468 (2007).

76. H. Hu, J. Gan, P. Jonas, Fast-spiking, parvalbumin+ GABAergic interneurons: From cellular design to microcircuit function. Science 345, 1255263 (2014).

77. R. G. Natan, W. Rao, M. N. Geffen, Cortical Interneurons Differentially Shape Frequency Tuning following Adaptation. Cell Reports 21, 878–890 (2017).

78. J. Urban-Ciecko, A. L. Barth, Somatostatin-expressing neurons in cortical networks. Nat Rev Neurosci 17, 401–409 (2016).

79. M. Okun, I. Lampl, Instantaneous correlation of excitation and inhibition during ongoing and sensory-evoked activities. Nat Neurosci 11, 535–537 (2008).

80. X.-J. Wang, Probabilistic Decision Making by Slow Reverberation in Cortical Circuits. Neuron 36, 955–968 (2002).

81. D. J. Amit, N. Brunel, Model of global spontaneous activity and local structured activity during delay periods in the cerebral cortex. Cereb Cortex 7, 237–252 (1997).

82. H. K. Inagaki, L. Fontolan, S. Romani, K. Svoboda, Discrete attractor dynamics underlies persistent activity in the frontal cortex. Nature 566, 212–217 (2019).

83. H. Sohn, D. Narain, Neural implementations of Bayesian inference. Current Opinion in Neurobiology 70, 121–129 (2021).

84. S. Sachidhanandam, V. Sreenivasan, A. Kyriakatos, Y. Kremer, C. C. H. Petersen, Membrane potential correlates of sensory perception in mouse barrel cortex. Nat Neurosci 16, 1671–1677 (2013).

85. C. C. H. Petersen, Sensorimotor processing in the rodent barrel cortex. Nat Rev Neurosci 20, 533–546 (2019).

86. A. R. Preston, H. Eichenbaum, Interplay of Hippocampus and Prefrontal Cortex in Memory. Current Biology 23, R764–R773 (2013).

87. J. Kim, A. Joshi, L. Frank, K. Ganguly, Cortical–hippocampal coupling during manifold exploration in motor cortex. Nature 613, 103–110 (2023).

88. M. Adibi, M. E. Diamond, E. Arabzadeh, Behavioral study of whisker-mediated vibration sensation in rats. PNAS 109, 971–976 (2012).

89. F. A. Wichmann, N. J. Hill, The psychometric function: I. Fitting, sampling, and goodness of fit. Perception & Psychophysics 63, 1293–1313 (2001).

90. M. Pachitariu, N. Steinmetz, S. Kadir, M. Carandini, H. K. D, Kilosort: realtime spike-sorting for extracellular electrophysiology with hundreds of channels. bioRxiv [Preprint] (2016). 10.1101/061481.

91. C. Rossant, S. N. Kadir, D. F. M. Goodman, J. Schulman, M. L. D. Hunter, A. B. Saleem, A. Grosmark, M. Belluscio, G. H. Denfield, A. S. Ecker, A. S. Tolias, S. Solomon, G. Buzsáki, M. Carandini, K. D. Harris, Spike sorting for large, dense electrode arrays. Nat Neurosci 19, 634–641 (2016).

92. D. Raposo, M. T. Kaufman, A. K. Churchland, A category-free neural population supports evolving demands during decision-making. Nat Neurosci 17, 1784–1792 (2014).

93. E. M. Meyers, D. J. Freedman, G. Kreiman, E. K. Miller, T. Poggio, Dynamic Population Coding of Category Information in Inferior Temporal and Prefrontal Cortex. Journal of Neurophysiology 100, 1407–1419 (2008).

94. V. Vapnik, “The Problem of Estimating Dependences from Empirical Data” in Estimation of Dependences Based on Empirical Data, V. Vapnik, Ed. (Springer, New York, NY, 2006; 10.1007/0-387-34239-7_1)*Information Science and Statistics*, pp. 1–26.

