## Supplementary Materials for "Cortical population codes for embedding sensory inputs into the prior context" (from another version of this preprint: version 1, 2026-03-02)

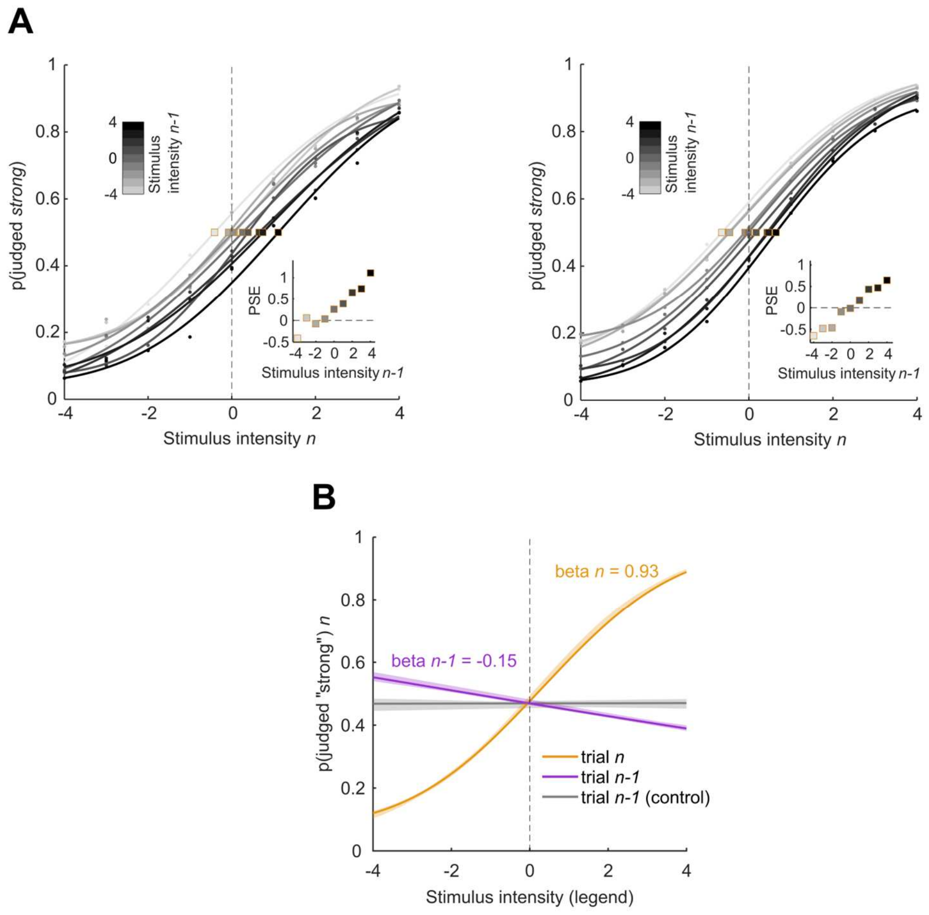


**Fig. S1. History-dependent perceptual decisions in rats during a vibrotactile psychophysical task.**

(**A**) Side-by-side comparison of behavioral data from the present study (left; ~ 18 000 trials collected during neural recordings), and data from (*5*) (right; ~ 100 000 trials collected in the absence of neurophysiological measurements). Of the 5 rats involved in this study, 3 rats were previously tested in Hachen et al. (2021). The plots represent psychometric curves for trial *n*, conditioned on *n-1* intensity. The squares denote the Point of Subjective Equality (PSE) for each curve. The inset represents the PSE as a function of *n-1* stimulus intensity. (**B**) The effect of stimulus *n-1* is reproduced as in Figure 1 (purple line), with the addition of a control analysis (grey line). To verify that the repulsive effect of *n-1* derived from perceptual decisions rather than spurious correlations across trials (see Methods), we simulated a dataset of choices from the trial *n* psychometric curve, thus excluding an effect of *n-1*, but with the same stimulus sequences as in the original dataset. In the presence of spurious correlations between *n-1* and *n* (e.g. overabundance of stimulus alternations), the multi parameter model could estimate a non-zero weight for *n-1* even in the absence of a real effect. In the data set derived from these simulated choices, the *n-1* effect was absent (gray line), supporting the interpretation that recent stimulus history – rather than stimulus ordering artifacts – drives the repulsive bias.

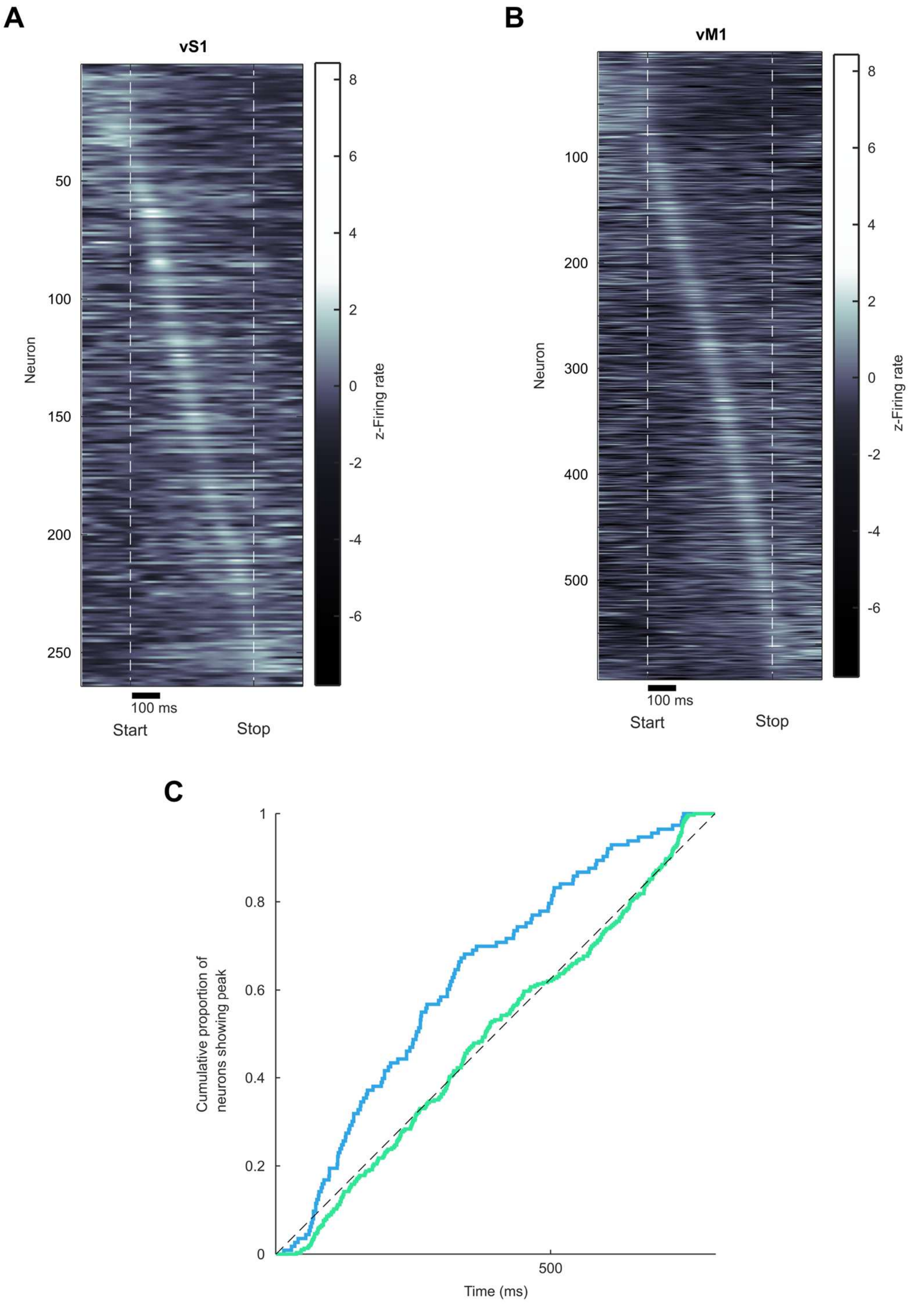


**Fig. S2. Individual-unit sensory responses in vS1 and vM1.**
(**A**) z-scored responses for all units in vS1. Units are further sorted by peak latency, in sequential order. Dashed lines mark stimulus onset and offset. (**B**) Same as (A) for all units in vM1. (**C**) Cumulative distribution of firing rate peaks for all neuronal units in the dataset, highlighting different response latencies in vS1 and vM1. The earliest timepoint considered (Time = 0) corresponds to stimulus onset.

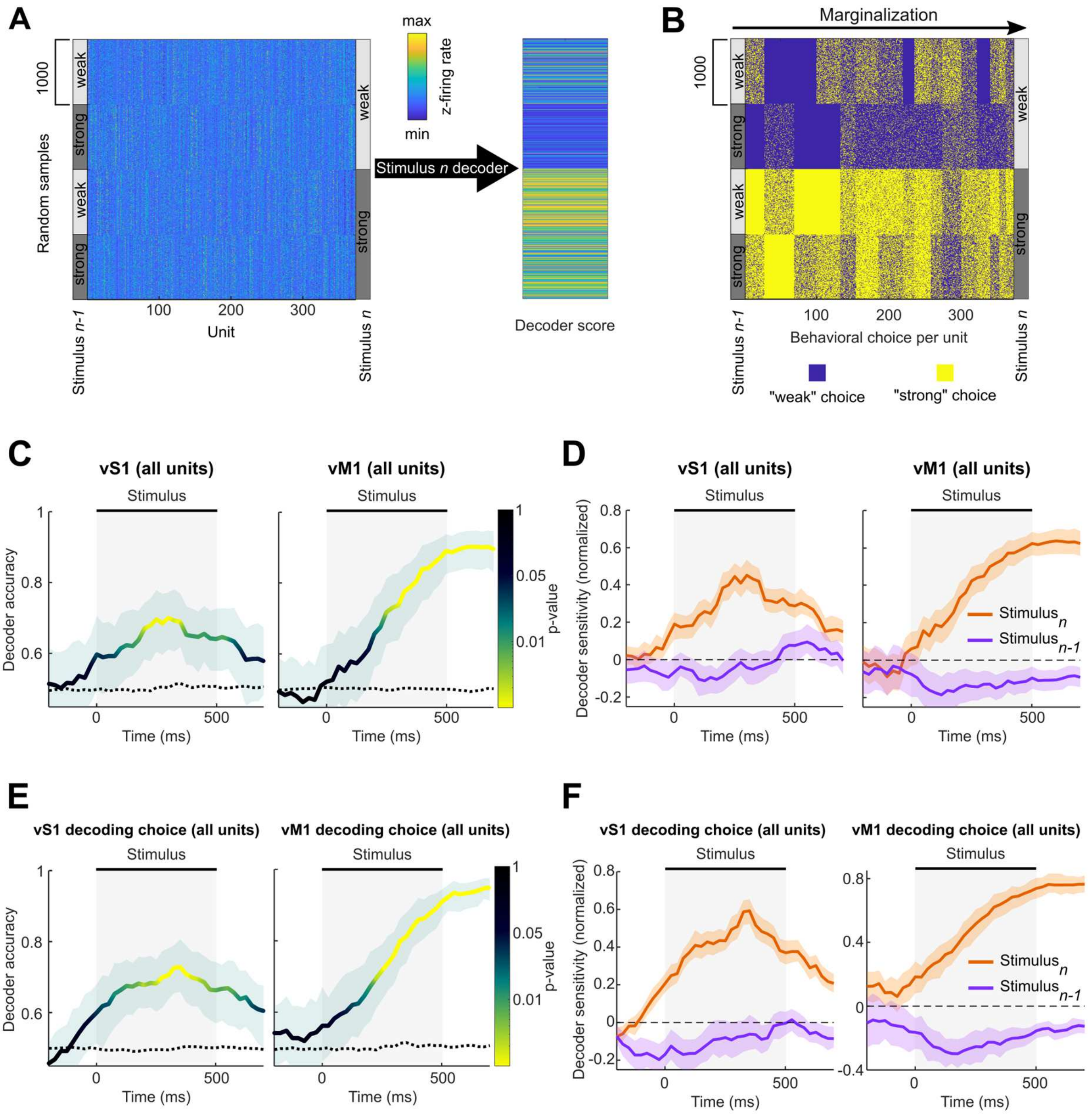


**Fig. S3. Population decoding of stimulus *n*.**

(**A**) Design of the population decoding analyses**.** Matrix resulting from the combination of resampled firing rates for each unit. The same amount of pseudo-simultaneous trials ("pseudo trials") was generated for each of the conditions of the 4x4 design matrix (in this example: Stimulus n x Stimulus n-1). For each cross-validation sample, two matrices of this kind were generated from different trials, one used for training and the other for testing the decoder. The classifier was trained to separate the pseudo trials according to only one of the two design conditions (in this case, Stimulus n). The distance of each pseudo trial from the boundary estimated by the classifier corresponds to the output "score" for that pseudo trial. Notice that, since the input to the classifier were z-scored firing rates, also the resulting "score" is expressed in the same unit. (**B**) Matrix storing the choice of the rat for each trial resampled from the units shown in a. Choice distributions were marginalized across columns, to yield the probability of judging stimulus n as "strong" for each pseudo-simultaneous population state. See Methods for further details. (**C**) Left: decoding accuracy of stimulus n from the firing rates of vS1 units in different time windows. Right: same for vM1 units. (**D**) Left: sensitivity of the vS1 stimulus n decoder to the value of stimulus n itself and to the value of stimulus n-1. Sensitivity was estimated for each time window included in the decoding analysis. For comparability between different analyses, all sensitivity values were normalized dividing all the values by their maximum. See main text and Methods for explanation of how sensitivity was computed. Right: Same for vM1 units. (**E**) Left: decoding accuracy of choice n from the firing rates of vS1 units in different time windows. Right: same vM1 units. (**F**) Left: sensitivity of the vS1 choice n decoder to the value of choice n itself and to the value of stimulus n-1. Sensitivity was estimated for each time window included in the decoding analysis. Right: same vM1 units. For comparability between different analyses, all sensitivity values were normalized dividing all the values by their maximum. See main text and Methods for explanation of how sensitivity was computed. Time windows had a 400 ms span and were separated by 25 ms. In all plots, abscissae denote the time point on which windows were centered.

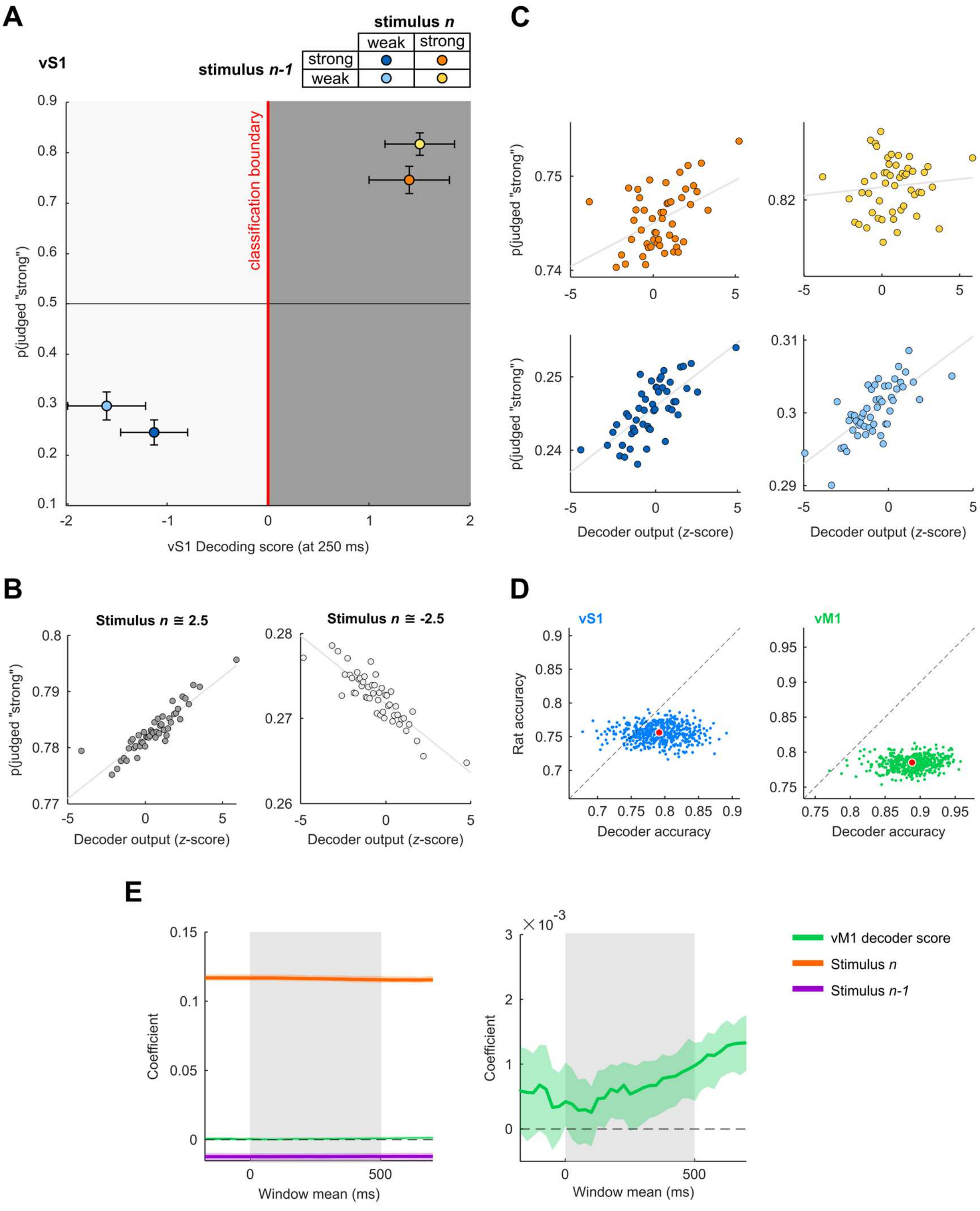


**Fig. S4. Relation between decoding scores and perceptual decisions.**

(**A**) Average cross-validated decoding scores and average rat judgments across 500 samples, for the 4 different trial types included in the analysis. Color codes for different categories of stimulus *n* and *n-1* are explained in the table at the top. Error bars in the main plots represent standard deviations. (**B**) Average rat judgment for each of the stimulus *n* categories as a function of decoding score for stimulus *n*. Scores were discretized in 50 equi-populated bins. To avoid potential confounds due to the presence of different values of speed within a single stimulus category, among all samples of trials we selected only those in which the mean stimulus (absolute) value was 2.5+-0.1 (6-7% of the decoded trials; see Methods). Gray lines correspond to regression slopes. (**C**) Average rat judgment within each of the 4 possible trial types represented in panel A (with the same color codes) as a function of decoding score for stimulus *n*. Scores were discretized in 50 equi-populated bins. To avoid potential confounds due to the presence of different values of speed within a single stimulus category, among all samples of trials we selected only those in which the mean stimulus (absolute) value was 2.5+-0.1 (6-7% of the decoded trials; Methods). Gray lines correspond to regression slopes. From top left clockwise, Pearson's rho = 0.67, 0.76, 0.92, 0.8; all p-values < 0.001), further controlling for the exact values of speed *n* and speed *n-1* within the sample (see caption). (**D**) Left: accuracy of rats and neural population decoding (at 250 ms) in the vS1 dataset. Right: accuracy of rats and population decoding (at 500 ms) in the vM1 dataset. Each point corresponds to a different cross-validated sample. (**E**) Coefficients estimated for the vM1-informed linear model of rat decisions (see main text). Lines represent means across cross-validation samples, error bars indicate standard errors of the mean. Left: full picture, right: zoomed-in picture.

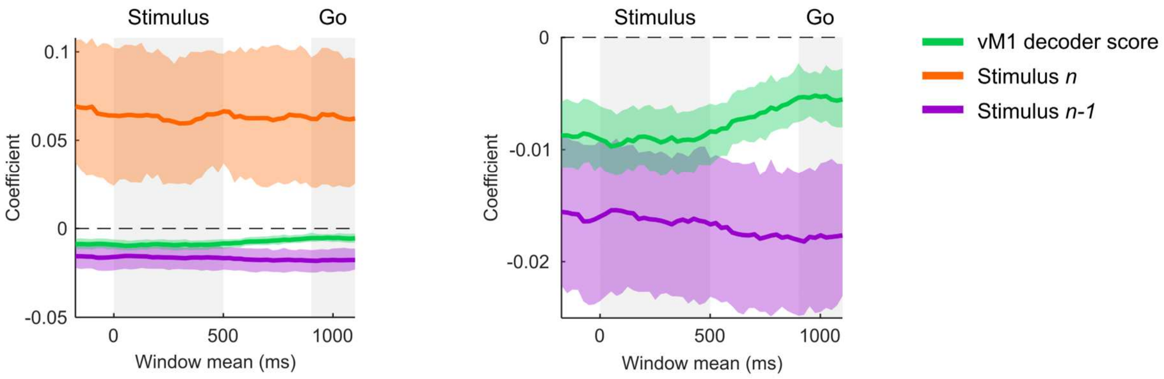


**Fig. S5. Population decoding of stimulus *n-1* and its relevance to behavioral choices**. Coefficients estimated for the vM1-informed linear model of rat decisions (see main text). Lines represent means across cross-validation samples, error bars indicate standard errors of the mean. Left: full picture, right: zoomed-in picture.

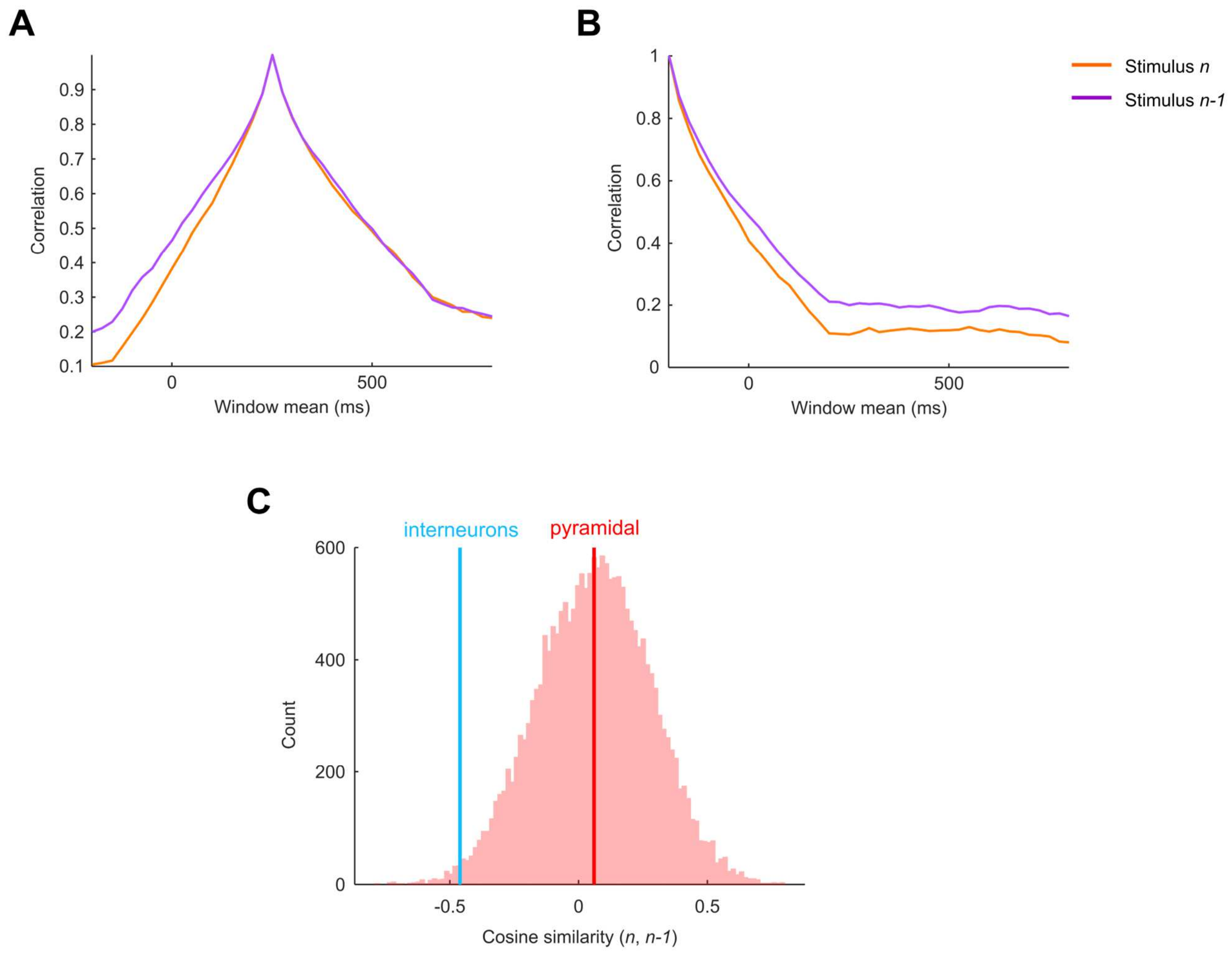


**Fig. S6. Distribution of population codes in vM1.**
(**A**) Correlation between the code estimated at 250 ms time (middle of vibration delivery) and the codes estimated at different times during the trial, for the n and the n-1 decoder. (**B**) Correlation between the code estimated at -200 ms time (middle of vibration delivery) and the codes estimated at different times during the trial, for the n and the n-1 decoder. (**C**) Cosine similarity values between the codes for stimulus n and n-1 obtained from the entire populations of pyramidal neurons and interneurons (vertical lines), together with histogram of cosine similarity values obtained by resampling pyramidal neurons.

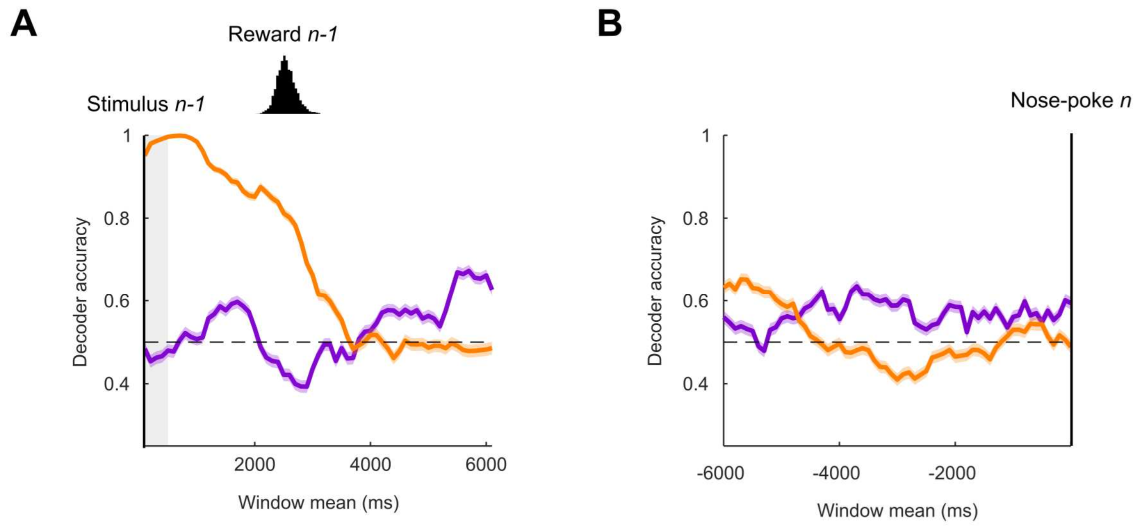


**Fig. S7. Transition between "real time" representation to "memory trace".**
(**A**) Cross-temporal validation of decoders trained to classify stimulus *n-1* using neural data sampled at two different timepoints: during stimulus *n-1* delivery (orange) or during stimulus *n* delivery (purple). Firing rates were aligned to stimulus onset (vertical line). The marginal histogram indicates reward delivery in trial n-1. (**B**) Cross-temporal validation of decoders trained to classify stimulus *n-1* using neural data sampled at two different timepoints: during stimulus *n-1* delivery (orange) or during stimulus *n* delivery (purple). Firing rates were aligned to trial n onset (nose-poke, vertical line).
